# Retrotransposon Activation Contributes to Neurodegeneration in a *Drosophila* TDP-43 Model of ALS

**DOI:** 10.1101/090175

**Authors:** Lisa Krug, Nabanita Chatterjee, Rebeca Borges-Monroy, Stephen Hearn, Wen-Wei Liao, Kathleen Morrill, Lisa Prazak, Yung-Heng Chang, Richard M Keegan, Nikolay Rozhkov, Delphine Theodorou, Molly Hammell, Josh Dubnau

**Affiliations:** Cold Spring Harbor Laboratory, Cold Spring Harbor, NY 11724, USA.; Watson School of Biological Sciences, Cold Spring Harbor Laboratory; Farmingdale State College, Farmingdale, NY 11735; Department of Anesthesiology, Stony Brook School of Medicine, 11794; Department of Neurobiology and Behavior, Stony Brook School of Medicine, 11794

## Abstract

Amyotrophic lateral sclerosis (ALS) and frontotemporal lobar degeneration (FTLD) are two incurable neurodegenerative disorders that exist on a symptomological spectrum and share both genetic underpinnings and pathophysiological hallmarks. Functional abnormality of TAR DNA-binding protein 43 (TDP-43), an aggregation-prone RNA and DNA binding protein, is observed in the vast majority of both familial and sporadic ALS cases and in ∼40% of FTLD cases, but the cascade of events leading to cell death are not understood. We have expressed human TDP-43 (hTDP-43) in *Drosophila* neurons and glia, a model that recapitulates many of the characteristics of TDP-43-linked human disease including protein aggregation pathology, locomotor impairment, and premature death. We report that such expression of hTDP-43 impairs small interfering RNA (siRNA) silencing, which is the major post-transcriptional mechanism of retrotransposable element (RTE) control in somatic tissue. This is accompanied by de-repression of a panel of both LINE and LTR families of RTEs, with somewhat different elements being active in response to hTDP-43 expression in glia versus neurons. hTDP-43 expression in glia causes an early and severe loss of control of a specific RTE, the endogenous retrovirus (ERV) *gypsy*. We demonstrate that *gypsy* causes the degenerative phenotypes in these flies because we are able to rescue the toxicity of glial hTDP-43 either by genetically blocking expression of this RTE or by pharmacologically inhibiting RTE reverse transcriptase activity. Moreover, we provide evidence that activation of DNA damage-mediated programmed cell death underlies both neuronal and glial hTDP-43 toxicity, consistent with RTE-mediated effects in both cell types. Our findings suggest a novel mechanism in which RTE activity contributes to neurodegeneration in TDP-43-mediated diseases such as ALS and FTLD.

**AUTHOR SUMMARY:** Functional abnormality of TAR DNA-binding protein 43 (TDP-43), an aggregation-prone RNA and DNA binding protein, is observed in the vast majority of both familial and sporadic ALS cases and in ∼40% of FTLD cases, and mutations in TDP-43 are causal in a subset of familial ALS cases. Although cytoplasmic inclusions of this mostly nuclear protein are a hallmark of the disease, the cascade of events leading to cell death are not understood. We demonstrate that expression of human TDP-43 (hTDP-43) in *Drosophila* neurons or glial cells, which results in toxic cytoplasmic accumulation of TDP-43, causes broad expression of retrotransposons. In the case of glial hTDP-43 expression, the endogenous retrovirus (ERV) gypsy causally contributes to degeneration because inhibiting gypsy genetically or pharmacologically is sufficient to rescue the phenotypic effects. Moreover, we demonstrate that activation of DNA damage-mediated programmed cell death underlies hTDP-43 and gypsy mediated toxicity. Finally, we find that hTDP-43 pathology impairs small interfering RNA silencing, which is an essential system that normally protects the genome from RTEs. These findings suggest a novel mechanism in which a storm of retrotransposon activation drives neurodegeneration in TDP-43 mediated diseases such as ALS and FTLD.

## INTRODUCTION

RTEs are “genomic parasites” – “selfish” genetic elements that are coded within our genomes and that replicate themselves via an RNA intermediate. After transcription, an RTE-encoded reverse transcriptase generates a cDNA copy, and this cDNA is inserted into a new genomic location at the site of double stranded DNA breaks created by an endonuclease activity encoded by the RTE (1). Unrestrained RTE activity has been demonstrated to be highly destructive to genomes, resulting in large-scale deletions and genomic rearrangements, insertional mutations, and accumulation of DNA double strand breaks (2). RTE-derived sequences constitute ∼40% of the human genome, a quantity which encompasses a surprisingly large number of functional RTE copies. Although multiple interleaved, highly conserved gene silencing systems have evolved to protect the genome by blocking RTE expression, certain RTEs are nevertheless expressed in some somatic tissues (3, 4) and can replicate in a narrow window during neural development, leading to *de novo* genomic insertions in adult brain (5-12). Moreover, a gradual deterioration of RTE suppression – and resultant increase in RTE activity – has been documented with advancing age in a variety of organisms and tissues (13-18), including the brain (19). One or more of the gene silencing systems that normally block genotoxic RTE expression may therefore be weakened with age. We advance the novel hypothesis that a broad and morbid loss of control of RTEs contributes to the cumulative degeneration observed with TDP-43 protein aggregation pathology that is observed in a variety of neurodegenerative disorders, including ALS and FTLD, and that this loss of control of RTEs is the result of a negative impact of TDP-43 pathology on general RTE suppression mechanisms that are most prevalently relied upon in somatic tissue such as the brain.

TDP-43 is a member of the hnRNP family that homodimerizes to bind single stranded RNA and DNA with UG/TG-rich motifs (20). This pleiotropic protein was originally identified as a transcriptional repressor that binds to the TAR element of the HIV-1 retrovirus to repress transcription (21). TDP-43 is capable of shuttling back and forth from the nucleus to the cytoplasm but is predominantly found in the nucleus in healthy cells. In cells that are experiencing TDP-43 protein pathology, the protein accumulates in dense cytoplasmic inclusions that include full-length protein, caspase cleavage products and C-terminal fragments of TDP-43, as well as abnormally phosphorylated and ubiquitinated protein (22-24). TDP-43 protein pathology is currently thought to involve toxicity incurred by cytoplasmic aggregates, interference with normal cytoplasmic function, depletion of normal nuclear TDP-43 stores, or some combination thereof (25).

Functional abnormality of TDP-43, an aggregation-prone RNA binding protein, is commonly observed in a spectrum of neurodegenerative diseases that spans motor neuron deterioration and progressive paralysis in ALS to dementia and cognitive decline in FTLD (26). 90% of ALS cases and a large fraction of FTLD cases are considered to be genetically sporadic, in the sense that no known genetic lesion precipitates pathology. However, loss of nuclear TDP-43 and accumulation of TDP-43 immunoreactive cytoplasmic inclusions is observed in nearly all ALS and almost half of FTLD cases (26-28). The mechanism that initiates the nucleation of TDP-43 protein pathology in apparently genetically normal individuals is not understood (27). However, TDP-43 contains a low complexity domain in its C-terminal region, which is a common feature of RNA binding proteins that exhibit aggregation pathology in a variety of neurodegenerative disorders. Indeed a recent literature has established that cellular stress can induce such low complexity domain proteins, including TDP-43, to undergo a concentration dependent phase separation to form liquid droplets that over time can drive fibrilization (29-31). TDP-43 protein also is known to accumulate in cytoplasmic stress granules in response to cellular stress (32). Importantly, nuclear TDP-43 protein normally regulates splicing of TDP-43 mRNA, leading to nonsense mediated decay of its own message (33). Thus the formation of cytoplasmic inclusions and clearance from the nuclear compartment that is observed in patients is also associated with loss of this feedback inhibition onto TDP-43 mRNA, leading to increased accumulation of cytoplasmic TDP-43 mRNA (34), which likely exacerbates formation of cytoplasmic inclusions.

Animal models of TDP-43 related disorders – and neurodegenerative disorders in general – have taken advantage of the concentration dependence of low complexity domain protein aggregation (35). Most animal models of neurodegenerative diseases therefore have involved transgenic expression to increase protein concentration above endogenous levels, and reproduce many of the signatures of human disease, which in the case of TDP-43 includes aggregation of TDP-43 protein in cytoplasmic inclusions and downstream neurological defects (26, 36, 37). Although such animal models are imperfect representations of what is largely a sporadically occurring disorder, they have enabled the delineation of a myriad of cellular roles for TDP-43 (26, 38) and have provided the means to uncover genetic interactions between TDP-43 and other genes that are implicated in neurodegenerative disorders (39-42). TDP-43 pathology in animal models is now understood to cause global alterations in mRNA stability and splicing, de-repression of cryptic splicing, and biogenesis of some microRNAs (miRNAs) (26, 27, 36, 37, 43-45). In principle, any of the cellular impacts of TDP-43 protein pathology could contribute to disease progression either alone or in combination. However, no clear consensus has yet emerged regarding the underlying causes of neurodegeneration in TDP-43 pathologies.

The RTE hypothesis investigated here is motivated by a series of prior observations. First, as mentioned above, LINE 1 RTEs are expressed in some somatic tissue (3, 4) and can actively replicate during normal brain development, leading to *de novo* genomic insertions in adult brain tissue (5-12), although the frequency of de novo insertions per cell is still hotly debated (46, 47). Second, increased RTE activity occurs in the brain during aging (19). Moreover, elevated expression of RTEs has been detected in a suite of neurodegenerative diseases (48-55) and reverse transcriptase biochemical activity of unknown origin has been shown to be present in both serum and cerebrospinal fluid (CSF) of HIV-negative ALS patients (56-59). More recently, a specific RTE, the human ERV HERV-K, was found to be expressed in post-mortem cortical tissue of ALS patients and their blood relatives (48, 53, 56) and transgenic expression of the HERV-K Envelope (ENV) protein in mice is sufficient to cause motor neuron toxicity (53). Finally, we have previously predicted via meta-analysis of RNA Immunoprecipitation (RIP) and Crosslinked RIP (CLIP) sequencing data that TDP-43 protein binds broadly to RTE-derived RNA transcripts in rodent and human brain tissue and that this binding is selectively lost in cortical tissue of FTLD patients (52). However to date, no studies exist which address whether TDP-43 pathology causes endogenous RTEs to become expressed *in vivo,* no reports have probed the functional impact of TDP-43 pathology on the natural mechanisms of RTE suppression employed by somatic tissue such as the brain, and no studies have investigated the toxic effects of endogenous RTE activation on nervous system function.

The destructive capacity of RTEs has been extensively documented in many other biological contexts, including a wealth of seminal data from *Drosophila melanogaster* (60-62). To test whether RTEs play a role in TDP-43 mediated neurodegeneration, we used an established *Drosophila* transgenic model which afforded the means to examine whether RTE activation causally contributes to TDP-43 mediated toxicity and cell death. We found that several hallmarks of TDP-43-induced degeneration are the result of activation of *the gypsy* ERV, an RTE that is structurally related to HERV-K, and that this activation leads to DNA damage mediated cell death. Moreover, we uncovered an inhibitory effect of TDP-43 expression on small interfering RNA (siRNA) mediated silencing, leading to broad activation of a panel of RTEs. These findings strongly suggest a broad impact of TDP-43 pathology on general RTE activity.

## RESULTS

### Neuronal or Glial Expression of hTDP-43 Induces RTE Expression

In order to determine whether RTEs contribute to TDP-43 pathological toxicity, we implemented an established animal model in which hTDP-43 is transgenically expressed in *Drosophila.* As with other animal models, including mouse, rat, fish, and *C elegans,* such expression reproduces many neuropathological hallmarks of human disease, likely via interference with endogenous protein(s) function (25, 36, 37, 63, 64). In *Drosophila,* there is an endogenous putative ortholog of TDP-43, TBPH. Null mutations in TBPH in flies are lethal (65), as is the case with mammalian TDP-43. Hypomorphic loss of TBPH results in neurodevelopmental defects as with the mammalian gene. Overexpression-mediated toxicity has formed the basis of the preponderance of studies on TDP-43 in animal models, and has revealed much of what is known regarding TDP-43 protein function and dysfunction, leading to the dominant hypotheses regarding mechanisms of pathogenesis wherein toxic cytoplasmic aggregates are thought to contribute to disease progression (25-27, 37, 38, 64, 66). To test the impact of expressing hTDP-43 on RTE expression, we first used RNA sequencing (RNA-seq) to profile transcript abundance. In patient tissue, TDP-43 protein pathology is observed in both neurons and glial cells (27) and an emerging literature has implicated glial cell toxicity in ALS (67-69). Toxicity of TDP-43 in glia has similarly been documented in animal models, including in *Drosophila* (70-73). We therefore examined the effects of transgenic hTDP-43 expression in the neuronal versus glial compartments of the brain.

We conducted paired-end Illumina based RNA-seq on head tissue of flies expressing either pan-neuronal (*ELAV* > hTDP-43) or pan-glial (*Repo* > hTDP-43) hTDP-43 compared with control flies that carried the hTDP-43 transgene alone with no Gal4 driver (hTDP-43 / +). We generated two independent sequencing libraries for each genotype from a population of animals that were 8-10 days post-eclosion. We generated a total of ∼900 million reads, or about 150 million reads per sample (Table S1), and conducted differential expression analysis (see methods). In order to identify effects both on gene transcripts and RTE transcripts (Fig 1A-1D; Table S2A-S3B), we included reads that map to repetitive elements using an analysis pipeline that we have previously reported (52, 74). Both glial (*Repo* > hTDP-43) and neuronal (*ELAV* > hTDP-43) expression of hTDP-43 caused differential expression of a number of cellular transcripts (Fig 1A and 1C; Table S2A and S3A) and transposons, most of which were RTEs or Class I elements (Fig 1B and 1D; Table S2B and S3B). In the case of differentially expressed genes, a broad spectrum of cellular processes were represented (see Table S2A and S3A), with both increases and decreases in expression level seen for many genes. This is broadly consistent with previously reported transcriptome analysis using tissue from ALS patients (75). In fact, the differentially expressed transcripts identified in our RNAseq experiments were significantly enriched for orthologs of genes that are implicated in ALS (ALS KEGG gene list; Fig. S1 and Table S2C). In contrast with differentially expressed genes, when examining transposon transcripts the majority of those that were differentially expressed exhibited elevated levels in response to hTDP-43 expression. This was particularly striking for glial TDP-43 expression (*Repo* > hTDP-43; Fig 1D), where 23 of 29 differentially expressed transposons showed higher levels relative to controls. The majority of the differential effects were observed in RTEs (Class I elements), although a few Class II elements were also represented (Fig 1B and 1D).

**Fig 1.**
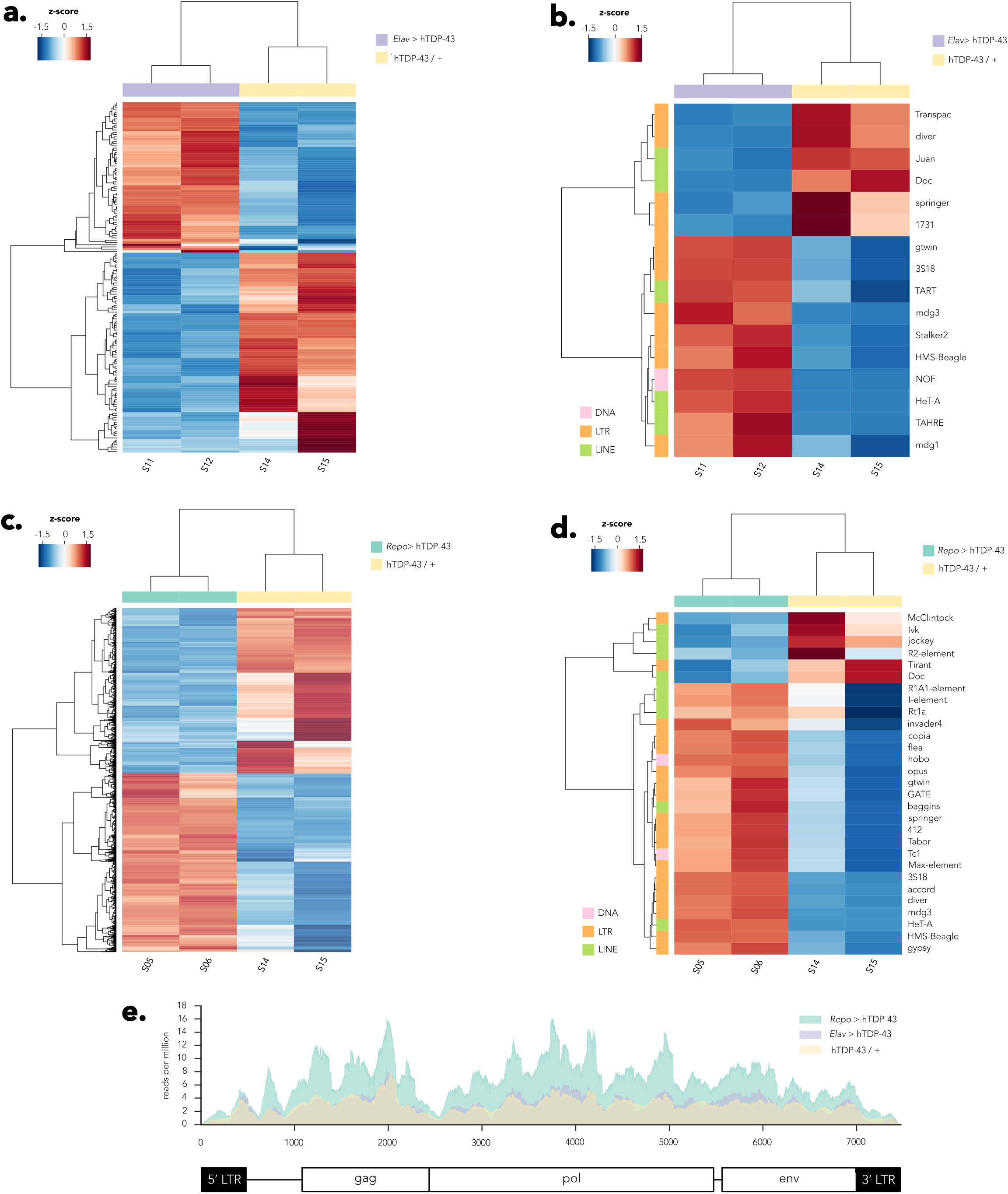
Neuronal and glial hTDP-43 expression results in induction of RTE expression. Differential expression of many genes and RTEs are detected in response to either neuronal or glial expression of hTDP-43 in head tissue of 8-10 day old flies (*N* = 2 biological replicates per genotype). (A) Neuronal *(Elav >* hTDP-43) expression of hTDP-43 results in both increases and decreases in expression of a broad variety of cellular transcripts (See Table S2A). (B) A panel of transposons, including many RTEs, also are impacted, with most exhibiting elevated expression (See Table S2B). (C) Glial expression of hTDP-43 *(Repo >* hTDP-43) also results in numerous transcriptome alterations, with many transcripts either increasing or decreasing in abundance (See Table S3A). (D) Many transposons, most of which are RTEs, exhibit elevated expression levels in response to glial hTDP-43 expression (See Table S3B). Several RTEs display elevated expression in response to both glial and neuronal hTDP-43 expression, however a number also exhibit specificity in response to either glial or neuronal hTDP-43 expression (compare Fig 1B and 1D). (E) The *gypsy* ERV exhibits elevated expression only in response to glial, but not neuronal, hTDP-43 expression. See methods for details regarding analysis pipeline, including statistical analysis.

It is also notable that while some RTE expression was elevated with both neuronal and glial hTDP-43 expression, there were several cases where effects were uniquely detected with only glial or only neuronal hTDP-43 expression. For example, the HeT-A LINE RTE and the mdg3, HMS-Beagle, gtwin and 3S18 LTR RTEs were elevated with either glial or neuronal expression of hTDP-43. However, the TART and TAHRE LINE RTEs and the Stalker2 and mdg1 LTR RTEs were only elevated in response to neuronal hTDP-43 expression, while a broad host of RTEs’ expression was elevated specifically in response to hTDP-43 expression in glia. Notable among these is the *gypsy* element, which we have previously demonstrated to be progressively de-repressed and even actively mobile with advanced age in brain tissue (19). We cannot formally rule out the possibility that some of the differences between differentially expressed RTEs in *Repo > hTDP-43* vs *Elav > hTDP-43* may result from variation in copy number of specific TEs between the two Gal4 strains. But we think this is unlikely to be a major contributing factor because all of the strains were backcrossed to the same wild type strain for a minimum of 5 generations prior to the experiments. “In the case *of gypsy,* expression levels are significantly increased in response to pan-glial hTDP-43 expression *(Repo >* hTDP-43) relative to controls (hTDP-43 / +) but no significant effect was observed with pan-neuronal expression of hTDP-43 (*ELAV* > hTDP-43) (Fig 1B, 1D, and 1E).

We selected the *gypsy* RTE as a candidate of interest to test the functional impact of loss of endogenous RTE suppression in response to hTDP-43 expression for several reasons. First, although *gypsy* was not the most abundantly expressed RTE in the RNA seq data, this element is known to be one of the most active natural transposons in *Drosophila melanogaster,* and is responsible for a high fraction of the spontaneous mutations that have been identified. Second, we have previously documented that *gypsy* is capable of replicating and generating *de novo* insertions in brain during advanced age (19). Third,*gypsy* is an ERV with functional similarity to HERV-K, which is expressed in some ALS patients (48, 53). And finally, because of intense prior investigation of the biology of this RTE, extant molecular genetic reagents provided the means to both perturb and detect gypsy function. We began by performing quantitative RT-PCR (qPCR) for both *ORF2 (Pol)* and *ORF3 (ENV)* of *gypsy* on head tissue of flies expressing either pan-neuronal (*ELAV* > hTDP-43) or pan-glial *(Repo >* hTDP-43) hTDP-43. Because disease risk is age dependent and symptoms in ALS patients are progressive, we also examined the compounding effects of age. At two relatively young ages (2-4 and 8-10 days post-eclosion) we observe a dramatic increase in expression of both ORFs (Fig 2A and 2B) of *gypsy* specifically in flies expressing hTDP-43 in glia. In contrast, flies expressing neuronal hTDP-43 experience a wave of *gypsy* expression at the population level that occurs much later in age (Fig S2A for *ORF3;* similar effects seen for *ORF2,* data not shown) in a similar manner to genetic controls that do not express hTDP-43 (see also: (19)). These flies do not show a significant impact of hTDP-43 expression on *gypsy* transcript levels. This is entirely consistent with the RNA-seq analyses (Fig 1B and 1D), where *gypsy* expression was found to be increased in head tissue specifically in response to glial hTDP-43 expression, but not to expression of hTDP-43 in neurons. Importantly, different genomic copy number or basal levels of *gypsy* expression between the parental Elav-Gal4 and Repo-Gal4 lines are unlikely to underlie the separate effects that we observe *on gypsy* when driving hTDP-43 expression in either neurons or glia (Figs. S2A.5 and S2A.6). Taken together, the RNA-seq and qPCR experiments confirm that *gypsy* RTE RNA levels are significantly and precociously elevated in response to pan-glial hTDP-43 expression.

**Fig 2.**
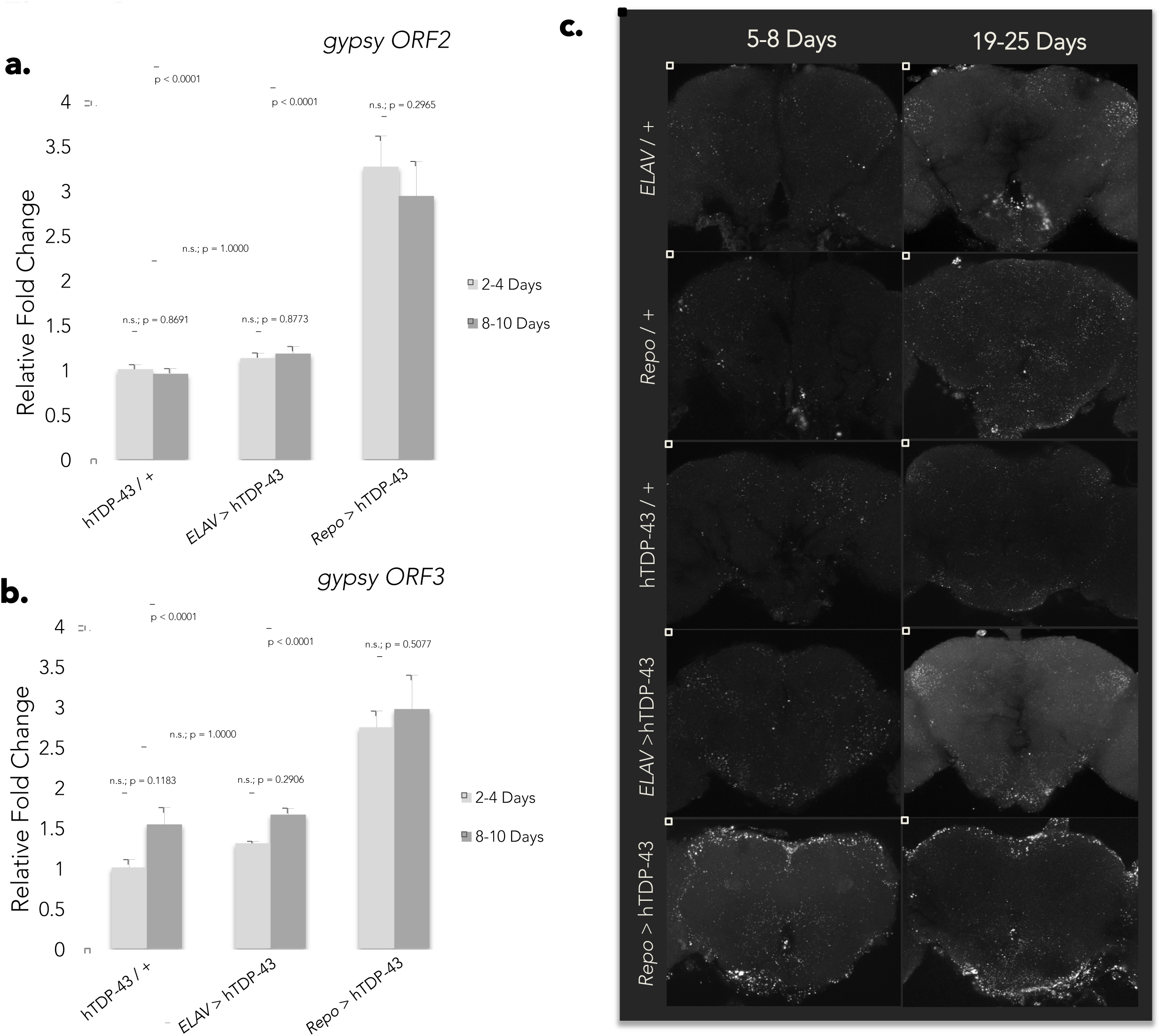
Glial hTDP-43 expression results in early and dramatic de-suppression of the gypsy ERV. (A) Transcript levels of *gypsy ORF2 (Pol)* as detected by qPCR in whole head tissue of flies expressing hTDP-43 in neurons (*ELAV* > hTDP-43) versus glia (*Repo* > hTDP-43) at a young (2-4 Day) or aged (8-10 Day) time point. Transcript levels normalized to *Actin* and displayed as fold change relative to flies carrying the hTDP-43 transgene with no Gal4 driver (hTDP-43 / +) at 2-4 Days (means + SEM). A two-way ANOVA reveals a significant effect of genotype (p < 0.0001) but no effect of age (p = 0.5414). *N =* 8 for all groups. (B) An equivalent analysis shows *that gypsy ORF3 (Env)* likewise displays a significant effect of genotype (p < 0.0001) and no effect of age (p = 0.6530). *N =* 4 for the 2-4 Day cohort and *N =* 5 for the 8-10 Day cohort. (C) Central projections of whole mount brains immunostained with a monoclonal antibody directed against *gypsy* ENV protein reveals dramatic, early accumulation of ENV immunoreactive puncta in brains expressing glial hTDP-43 (5-8 Days) in comparison to both age-matched genetic controls (*ELAV*/ + ; *Repo /* + ; hTDP-43 / +) and flies expressing neuronal hTDP-43. This effect persists out to 19-25 Days post-eclosion. *ELAV/* +, 5-8 Day (*N=* 3), 19-25 Day (*N =* 4); *Repo /* +, 5-8 Day (*N =* 3), 19- 25 Day (*N =* 3); hTDP-43 / +, 5-8 Day (*N =* 5), 19-25 Day (*N =* 2); *ELAV>* hTDP-43, 5-8 Day (*N =* 2), 19-25 Day (*N* =4); *Repo >* hTDP-43, 5-8 Day (*N* = 7), 19-25 Day (*N = 8).*

Whole mount immunolabeling of brains using a monoclonal antibody directed against the *gypsy* ENV glycoprotein (19, 76) likewise shows early (5-8 days post-eclosion) and acute accumulation of strongly immunoreactive puncta particularly in brains of flies expressing glial hTDP-43 (Fig 2C; for quantification see Fig S2B). These intense puncta are observed throughout the superficial regions, which contain the majority of cell somata, as well as in deeper neuropil (Fig 2C and data not shown) and persist into older ages. In contrast, we do not observe neuronal hTDP-43 expression to cause elevated *gypsy* levels above that seen in wild type flies at any time point with either qPCR or immunolabeling (Fig 2C and Fig S2A and data not shown). Given that effects of glial hTDP-43 expression on *gypsy* ENV immunoreactivity were so robust in 5-8 day old animals, we examined ENV at earlier time points. We found that in animals expressing hTDP-43 in glia, there is little detectable *gypsy* ENV protein expression at 0 days (immediately following eclosion). In brains from animals 3 days post eclosion, we observe regional puncta with a variable intensity and spatial location (Fig S2C) although this effect was difficult to quantify because of its variability.

We have previously demonstrated that in brains of very old animals, *gypsy* is both expressed and actively mobile, leading to accumulation of *de novo* insertions. Given the abnormally high levels of *gypsy* expression that we observed in brains of young animals in response to hTDP-43 expression in glial cells, we wondered whether *gypsy* was also actively mobile in this context. To test this, we designed a gypsy reporter construct that turns on a nuclear mCherry reporter after only after replicating through an RNA intermediate and reinserting into the genome (Fig. S3A). This strategy has been successfully used to report mammalian LINE element mobilization(77). We found that this reporter detects an age-dependent increase in *gypsy* retrotransposition in wild type brains (Fig. S3B,C) as we previously reported(19). This effect is significantly exacerbated when hTDP-43 is expressed in glial cells (Fig. S3B and S3C).

### Neuronal and Glial hTDP-43 Expression Causes Age-Dependent Neurological Deterioration

We next examined the relative impact of glial and neuronal hTDP-43 expression on the physiological health of the animal. As previously documented (71-73), we see effects with either neuronal or glial expression. However we observe differing severity and time courses, with effects of glial expression being more acute than those observed with expression in neurons. Flies expressing hTDP-43 in neurons exhibit significant locomotor impairment at 1-5 days post-eclosion, and flies expressing glial hTDP-43 show more severe locomotor impairment at this same age. This effect is further exacerbated by 5-10 days post-eclosion; at which point the animals expressing hTDP-43 in glia are largely immobile (Fig 3A). As previously reported (66, 71, 72, 78-81), flies expressing neuronal hTDP-43 exhibit reduced lifespan in comparison to genetic controls. But flies expressing hTDP-43 in glia display an even more severely reduced lifespan (Fig 3B). We further observe rampant cell death as detected by terminal deoxynucleotidyl transferase dUTP nick end labeling (TUNEL) in the brains of flies expressing hTDP-43 in glia as early as 5 days post-eclosion (Fig 3C). Similarly, with transmission electron microscopy (TEM; Fig 3D) we observe profuse proapoptotic nuclei in brains of 12 day-old flies expressing glial hTDP-43. In contrast, driving expression of hTDP-43 in neurons under the *OK107-Gal4* driver, which provides high levels of expression in the well-defined and easily imaged population of central nervous system (CNS) neurons that constitute the mushroom body, results in little to no increase in TUNEL labeling (consistent with a previous report: (80)) even when the flies were aged to 30 days (Fig S4A). The relative expression of hTDP-43 under the two major Gal4 drivers we are using, *Repo-Gal4* (glia) and *ELAV-Gal4* (neurons), does not differ with age, suggesting that divergent age effects on expression level cannot account for the observed differences in toxicity and impact on physical health (Fig S4D and S4E; respectively). Furthermore, we do not observe any effect of hTDP-43 expression on levels of the endogenous fly ortholog, *TBPH,* regardless of cell type of expression (Fig S4F; Table S4). Thus, the phenotypes that we observe are not caused by indirect effects on *TBPH* transcript abundance but instead derive from toxicity of the hTDP-43 transgene itself. The levels of expression of the hTDP-43 transgene relative to the endogenous TBPH gene also are similar to what has been reported in rodent models (Table S4). As is true in other animal models and in human patients, we cannot readily distinguish whether the effects we observe are due to toxic gain of function, dominant interference with an endogenous protein, or some combination thereof. Importantly, however, we can detect a disease specific phosphorylated isoform of hTDP-43 (Fig S4B) as well as cytoplasmic accumulation and nuclear clearance of the protein (Fig S4C), implying that the human protein is being processed in the CNS of the fly as it is thought to be in the disease state in human tissue.

**Fig 3.**
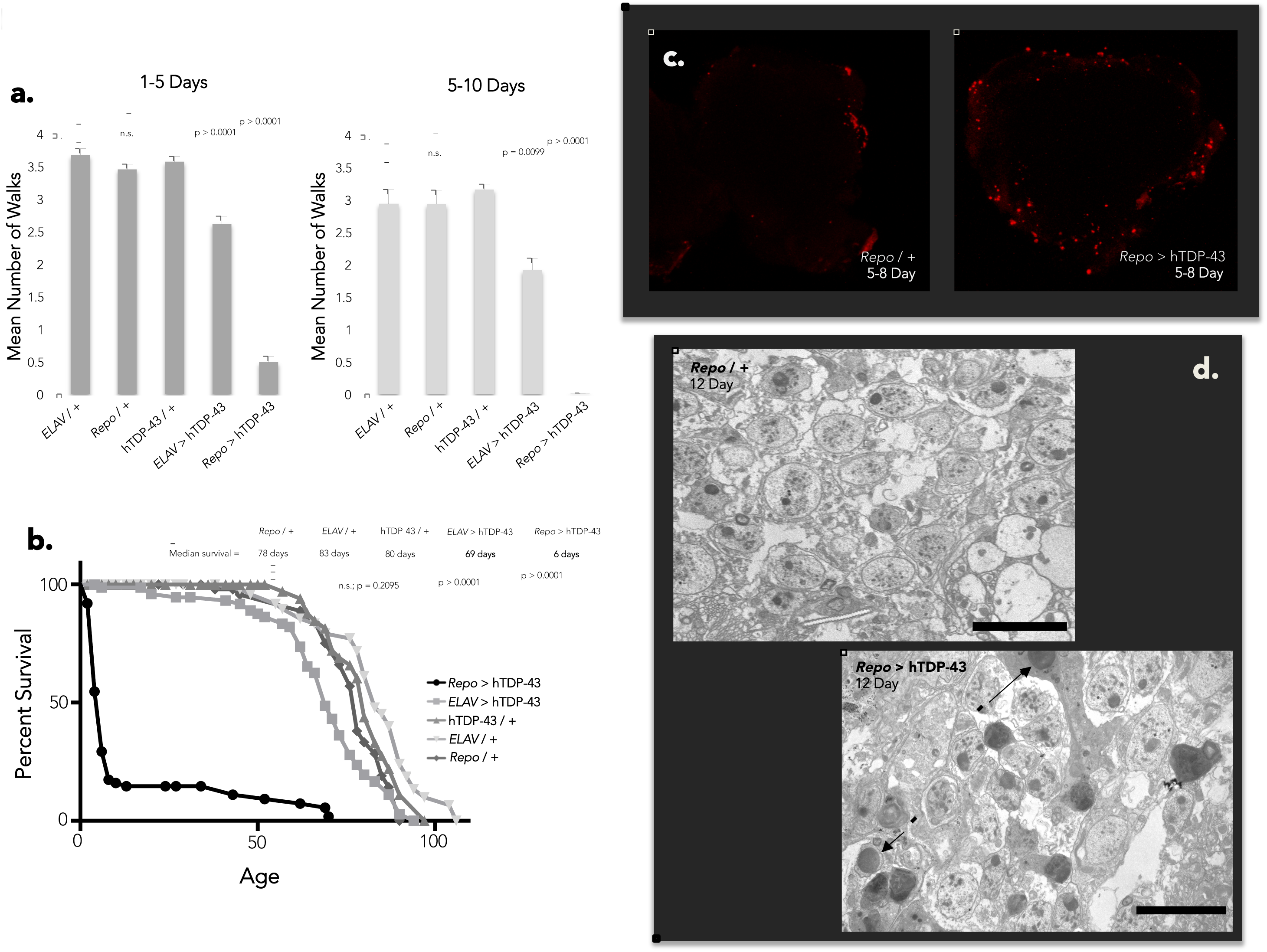
Neuronal and glial hTDP-43 expression induces physiological impairment and toxicity with varying severity. (A) Flies expressing glial hTDP-43 display extreme locomotor impairment at 1-5 days post-eclosion in the Benzer fast phototaxis assay, while flies expressing neuronal hTDP-43 demonstrate a slight locomotor deficit in comparison to genetic controls (one-way ANOVA, p < 0.0001). This trend continues and is exacerbated by 5-10 days post-eclosion (one-way ANOVA, p < 0.0001). Four biological replicates performed for each experiment. (B) Lifespan analysis of flies expressing neuronal versus glial hTDP-43 in comparison to genetic controls. (C) Central projections of whole-mount brains reveals a stark increase in TUNEL-positive cells in flies expressing glial hTDP-43 in comparison to genetic controls at 5 days post-eclosion. *N =* 16 for *Repo /* + and *N =* 18 for *Repo >* hTDP-43. (D) TEM likewise reveals rampant apoptosis in the neuropil of flies expressing glial hTDP-43 at 12 days post-eclosion. Arrowheads indicate pro-apoptotic nuclei, as identified by morphology.

### RTE and Chk2 Activity Mediate Effects of hTDP-43 on Lifespan and Cell Death

Our observation that a panel of RTEs are expressed in response to hTDP-43 transgene expression, along with the extensively documented toxic effects of loss of control of RTEs in other biological contexts (19, 60-62) and our observation that the *gypsy* RTE itself can actively replicate and generate *de novo* insertional mutations in brain tissue during aging (19), and in response to the hTDP-43 transgene, suggested the possibility that loss of *gypsy* silencing might in fact account for a portion of the physiological toxicity observed with hTDP-43 expression in glia. To test whether the *Drosophila* ERV *gypsy* causally contributes to the harmful effects of hTDP-43, we used a previously published inverted repeat (IR) “RNAi” construct (55) directed against *gypsy ORF2 (gypsy(IR))* that is sufficient to reduce the expression of *gypsy* by approximately 50% in head tissue of 28-day old animals (Fig S5A). We found that co-expression of this *gypsy* (IR) substantially ameliorates the lifespan deficit induced by glial hTDP-43 expression (Fig 4A). This effect is not observed when a control IR construct is co-expressed with hTDP-43 in glial cells (*Repo* > hTDP-43 + GFP(IR); Fig 4B), and neither *the gypsy(IR)* nor the GFP(IR) constructs, when expressed alone under *Repo-Gal4* (Fig S5B) or *ELAV-Gal4* (Fig S5C) or when present without a Gal4 driver (Fig S5D), has such an effect on lifespan. Therefore activation of *gypsy* is responsible for a substantial portion of the toxicity that we observe when hTDP-43 is expressed in glia, which results in drastically premature death in these animals. In contrast, co-expression of *gypsy*(IR) does not rescue the lifespan deficit exhibited by animals expressing hTDP-43 in neurons (Fig 4C). This is in accordance with our observations from RNA-seq (Fig 1B and 1D), qPCR (Fig 2A and 2B; Fig S2A.1-S2A.3) and immunolabeling (Fig 2C) that neuronal expression of hTDP-43 also does not elevate *gypsy* expression above wild type levels at any given time point over the course of lifespan. The glial specificity *of gypsy(IR)* lifespan rescue is consistent with our observation that *gypsy* expression is induced specifically when TDP-43 is expressed in glia, lending credence to the conclusion that *gypsy* is causally participating in the resulting degenerative phenotype We of course cannot rule out the possibility that the *gypsy-*RNAi construct may also impact *gypsy* family RTEs that share sequence homology to *gypsy.* The conclusion that RTEs contribute to TDP-43 toxicity is further supported by the mild but significant lifespan extension that we observe with pharmacological inhibition of the reverse transcriptase activity that is essential for all RTE replication (Fig. S5E,F,G and H).

**Fig 4.**
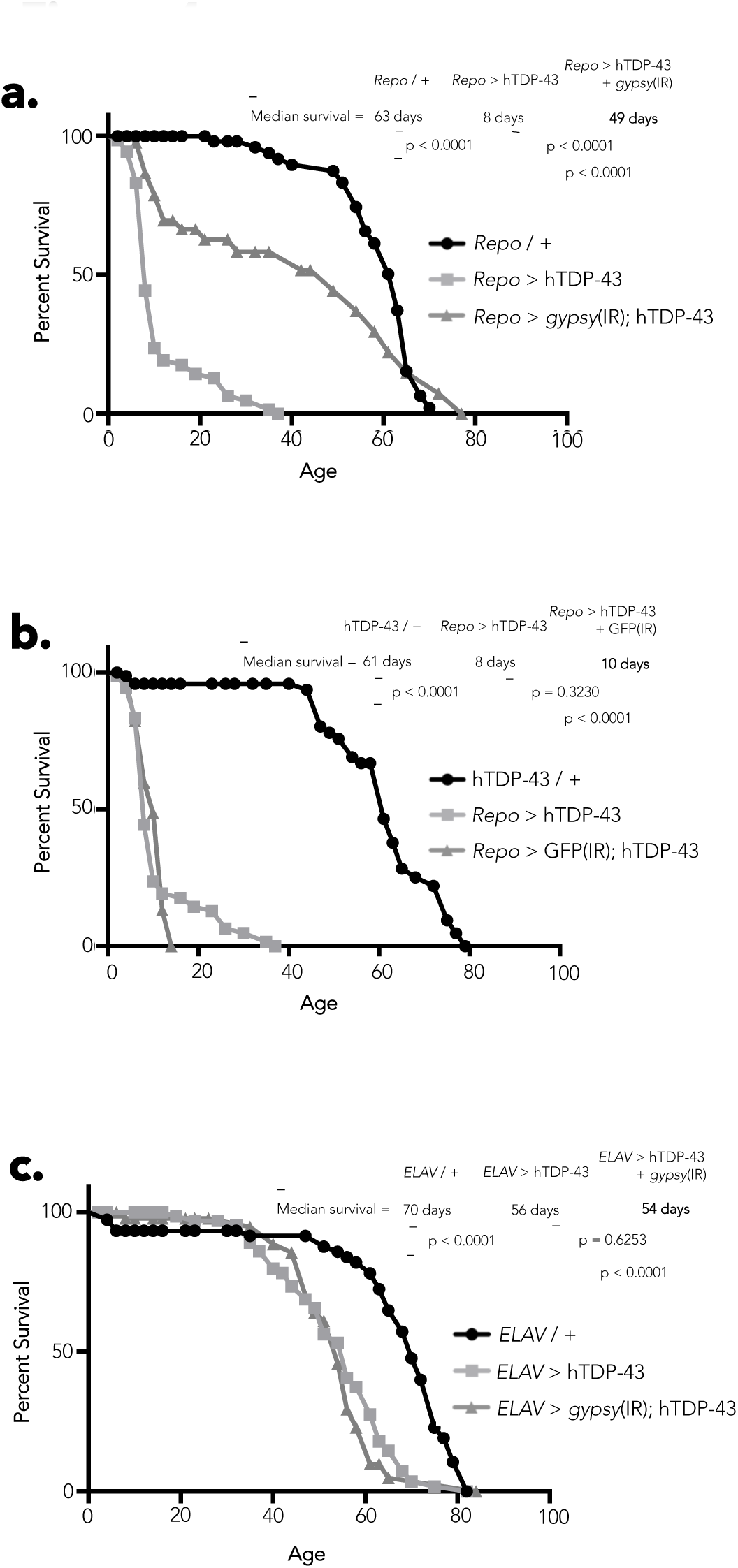
*gypsy* ERV expression contributes to hTDP-43 mediated toxicity. (A) Lifespan analysis shows that co-expression *ofgypsy(IR) (Repo > gypsy(IR)* + hTDP-43) partially rescues the lifespan deficit exhibited by flies expressing glial hTDP-43 *(Repo >* hTDP-43). (B) Co-expression of an unrelated GFP(IR) control transgene *(Repo >* GFP(IR) + hTDP-43) does not effect the lifespan of flies expressing glial hTDP-43 *(Repo >* hTDP-43). (C) Co-expression *of gypsy(IR) (ELAV > gypsy (IR) +* hTDP-43) has no effect on lifespan in flies expressing neuronal hTDP-43 *(ELAV>* hTDP-43).

RTE replication involves reverse transcription, generation of chromosomal DNA breaks, and integration of the RTE cDNA copy. DNA damage is therefore associated with either abortive or successful attempts at RTE replication and such DNA damage is thought to be a major source of cellular toxicity caused by RTE activity because it activates Chk2 signaling, which leads to programmed cell death. To test whether the harmful effects of hTDP-43 are in fact mediated by DNA damage, we capitalized on the previously documented ability of mutations in Chk2 to mask the toxic effects of RTE-induced DNA damage (82, 83). Importantly, mutations in Chk2 do not prevent accumulation of DNA damage; rather they prevent the signaling required for the cell to recognize that DNA damage has occurred and respond by committing to a programmed cell death pathway (84). We therefore employed an IR construct directed against *loki* (loki(IR)), the *Drosophila* ortholog of *chk2,* which is sufficient to significantly reduce levels of endogenous *loki* mRNA in head tissue of 28-day old animals (Fig S6A). Remarkably, co-expression of loki(IR) with hTDP-43 is able to fully rescue the lifespan deficit caused by hTDP-43 expression in glia (Fig 5A) or neurons (Fig 5B). These findings support the conclusion that DNA damage makes a major contribution to the loss of lifespan induced by either neuronal or glial expression of hTDP-43. This conclusion is supported by our RNA-seq findings, in which we observe that in each case the expression of a panel of RTEs is activated. Although neuronal hTDP-43 expression does not impact the levels of the gypsy RTE specifically, several other RTEs exhibit elevated expression (see: Fig 1B). Importantly, this extension of lifespan is not seen with co-expression of a control IR construct *(Repo >* hTDP-43 + GFP(IR); Fig 4B), and neither the GFP(IR) nor *loki(IR)* constructs when expressed individually under *Repo-Gal4* or *ELAV-Gal4* or present without a Gal4 driver (Fig S5B – S5D), have such an effect on lifespan on their own. These data suggest that Loki/Chk2 activity makes a major contribution to the pathological toxicity of hTDP-43 that we observe with both glial and neuronal hTDP-43 expression.

**Fig 5.**
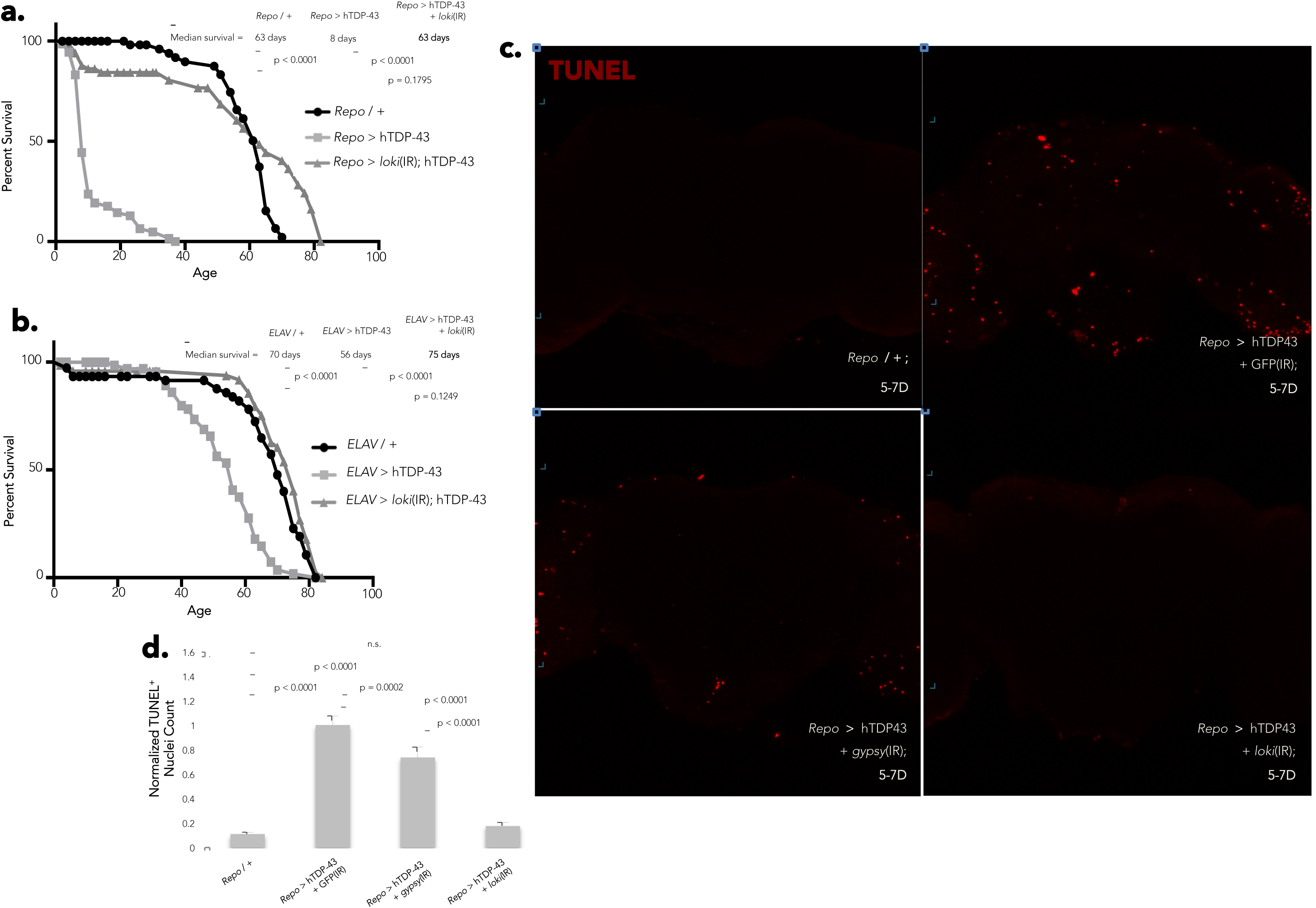
DNA damage-induced cell death and *gypsy* ERV expression contribute hTDP-43 mediated toxicity. (A) Lifespan analysis shows that co-expression *ofloki(IR) (Repo > loki(IR)* + hTDP-43) fully rescues the lifespan deficit exhibited by flies expressing glial hTDP-43 *(Repo >* hTDP-43). (B) Co-expression ofloki(IR) *(ELAV> loki(IR) +* hTDP-43) likewise fully rescues the lifespan deficit exhibited by flies expressing neuronal hTDP-43 *(ELAV>* hTDP-43). (C) Central projections of whole-mount TUNEL stained brains reveal a noticeable reduction in the apoptotic activity induced by glial hTDP-43 expression *(Repo >* hTDP-43 + GFP(IR)) *when gypsy* expression is knocked down *(Repo >* hTDP-43 *+ gypsy(IR)),* while knocking down *loki* completely alleviates the apoptosis induced by glial hTDP-43 expression *(Repo >* hTDP-43 + loki(IR)). (D) Quantification of (H), normalized to the positive control *(Repo >* hTDP-43 + GFP(IR)). *N =* 12 for *Repo /+;N=9* for *Repo >* hTDP-43 + GFP(IR); *N = 7* for *Repo >* hTDP-43 + *gypsy(IR);* and *N = 7* for *Repo >* hTDP-43 + loki(IR). *All of the lifespans with the exception of the NRTI feeding experiments shown in Fig 4, 5, and S3 were performed concurrently in order to ensure comparability across groups. Therefore, appropriate controls are shared across panels.

The brains of flies expressing hTDP-43 in glia display rampant cell death, seen both with TUNEL staining (Fig 3C) and at the level of TEM (Fig 3D). To test whether the decision of cells to commit to a programmed cell death pathway in response to hTDP-43 expression is mediated by Loki (Chk2), we co-expressed the *loki(IR)* that was so effective in suppressing hTDP-43 toxicity in survival analyses *(Repo >* hTDP-43 + loki(IR)) and found that this was sufficient to abolish the dramatic accumulation of TUNEL-positive nuclei induced by glial expression of hTDP-43 (Fig 5C and 5D). Moreover, we found that the *gypsy* RTE contributes at least in part to the decision of cells to undergo programmed cell death in response to hTDP-43 expression in glia, as knocking down gypsy (*Repo* > hTDP-43 + *gypsy*(IR)) also significantly reduces the TUNEL labeling observed in the CNS of these animals (Fig 5C and 5D). These effects are specific to *loki(IR)* and gypsy(IR) as co-expression of an unrelated UAS-(IR) construct with hTDP-43 in glia *(Repo >* hTDP-43 + GFP(IR)) does not significantly alter the number of TUNEL positive cells compared to brains of flies expressing hTDP-43 alone under *Repo-Gal4* (Fig S6B). Importantly, co-expression of the GFP(IR), *loki*(IR), and gypsy(IR) constructs with hTDP-43 under *Repo-Gal4* also does not significantly reduce the expression of hTDP-43 *(TARDBP;* Fig S6C). Differences in the level of hTDP-43 expression between these experimental groups therefore cannot account for the phenotypic rescue observed with *loki* or *gypsy* knock down in either the survival or cell death assays. Taken together, these data support the conclusion that the cell death induced by hTDP-43 is mediated predominantly via Loki/Chk2 activity in response to DNA damage, and that this DNA damage is likely induced by RTE activity. For both the physiological toxicity and cell death induced by hTDP-43 expression in glial cells, this effect is in large part due to the activity of one particular RTE, the *gypsy* ERV. These observations are in agreement with the well-documented accumulation of DNA double strand breaks induced by unleashing RTEs (85), as well as reports that transgenic expression of the HERV-K ENV protein in mice results in loss of volume in the motor cortex and DNA damage (53). While the impact *of gypsy* appears to be restricted to the case where hTDP-43 is expressed in glial cells, our RNA-seq data demonstrate that expression of hTDP-43 causes the induction of a panel of RTEs that normally would be silenced. Such results lead us to postulate that hTDP-43 pathology might be impacting the natural mechanisms by which RTEs in general are normally kept suppressed. We therefore designed a reporter assay to detect the effect of hTDP-43 expression on the siRNA system, which provides the primary silencing mechanism to keep RTEs in check in somatic tissues such as the brain.

### Expression of hTDP-43 Disrupts siRNA-Mediated Silencing

The major post-transcriptional RTE silencing system available in somatic tissue such as the brain is the siRNA pathway (86-91). siRNAs with sequence complementarity to RTEs have been detected in many species, including mammals (1, 87, 92), and RTE-siRNA levels have been demonstrated to affect RTE activity (1, 93-95). Moreover, disruptions in the siRNA pathway result in increased TE transcript levels (19, 90, 96) as well as novel insertions in the genome (19, 97). Indeed, we have previously shown that disruption of the major siRNA pathway effector *Argonaute 2 (Ago2)* leads to precocious *gypsy* expression in *Drosophila* head tissue and this is accompanied by rapid age-dependent neurophysiological decline (19). We therefore engineered a genetically encoded sensor system to inform us as to whether hTDP-43 expression impairs the efficiency of Dicer-2 (Dcr-2)/Ago2-mediated siRNA silencing in the *Drosophila* nervous system *in vivo.*

Our reporter system relied on three components. We co-expressed a Dcr-2 processed IR construct directed against GFP (GFP(IR)) with a GFP transgenic reporter. By selecting an effective GFP(IR), we were able to generate substantial silencing of the GFP reporter (Fig 6A and 6B). To test the effects of hTDP-43 on siRNA mediated silencing, we then co-expressed our third component: either hTDP-43 or an unrelated control transgene (tdTomato). This tripartite system was expressed either in all glial cells using the *Repo-Gal4* driver (Fig 6A) or in mushroom body neurons using the *OK107-Gal4* driver (Fig 6B). Brains of young (2-4 day) and middle aged (10-12 days) flies were imaged using confocal microscopy. In the case of neuronal expression we were able to carry the experiment out to old age (45-47 days), but this was not possible with glial expression of hTDP-43 as it results in dramatic reduction in lifespan (see Fig 3B). What we observed was conspicuously reminiscent of hTDP-43’s impact on *gypsy* expression. Glial expression of hTDP-43 causes a marked reduction of siRNA silencing efficacy, resulting in easily detectable expression of the GFP reporter. Such expression is dramatic and significant in brains of 2-4 day old flies and persists out to 10-12 days of age (Fig 6A). Brains are obviously deteriorated by the 10-12 day time-point (data not shown), which likely explains why GFP levels appear to drop off somewhat. Neuronal expression of hTDP-43 in the mushroom body has a similar but more slowly progressing effect on siRNA-mediated silencing of our GFP reporter, with a somewhat later onset (Fig 6B). Indeed, when we perform an analogous experiment using an endogenous reporter of siRNA mediated silencing in a separate structure we observe a similar effect. The *GMR-Gal4* driver, which drives high levels of expression in the fly eye, was used to express an IR construct directed against the endogenous *white*^*+*^ pigment gene in place of GFP as a reporter (Fig 6C). As with mushroom body neurons in the CNS, expression of hTDP-43 in the eye causes a progressive de-repression of the silenced reporter. It is noteworthy that the erosion of siRNA efficacy caused by hTDP-43 expression in the eye manifests as clusters of red-pigmented cells, a phenotype which is evocative of the stochastic clusters of ENV immunoreactivity observed early in response to glial hTDP-43 expression (Fig 6C and Fig S2C). In contrast, simply turning on expression of *white*^*+*^ after development results in a uniform darkening of the eye with age (Fig S7A and S7B). Taken together, these findings demonstrate that hTDP-43 expression interferes with siRNA-mediated silencing in several tissue types, resulting in de-suppression of reporter expression. In neurons hTDP-43 expression causes age-dependent progressive erosion of siRNA efficacy, while glial expression of hTDP-43 results in more acute siRNA silencing impairment.

**Fig 6.**
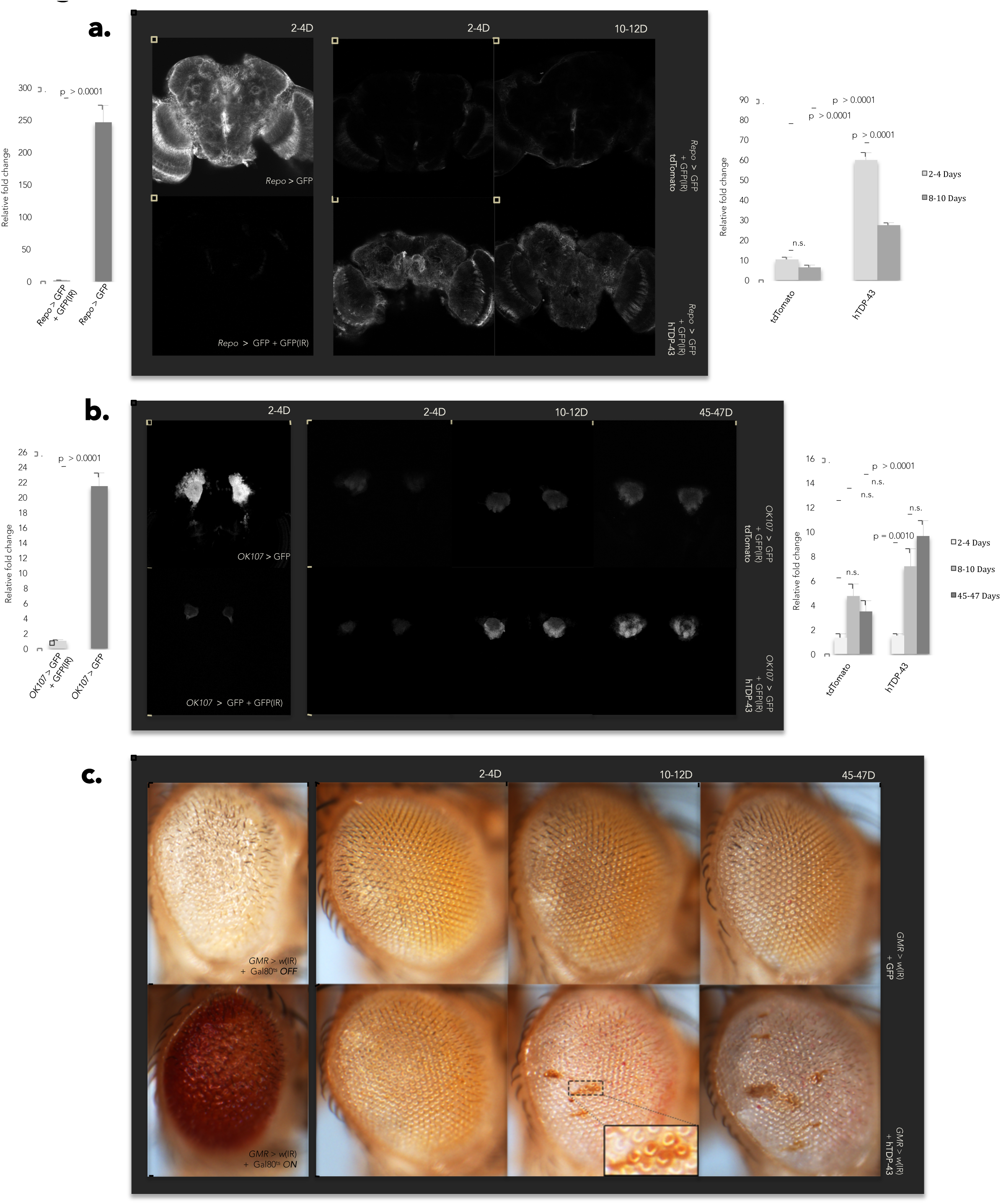
Glial and neuronal hTDP-43 expression erodes siRNA-mediated silencing. (A) Representative central projections show that co-expression of the hTDP-43 transgene, but not an unrelated tdTomato control transgene, interferes with the ability of a Dcr-2 processed IR (GFP(IR)) to silence a GFP transgenic reporter in glial cells using the *Repo-GAL4* driver. Quantification of GFP signal for each group is shown in the appropriate bar graph; values are represented as relative fold change over *Repo >* GFP + GFP(IR) (mean + SEM). A two-way ANOVA reveals significant effects of both genotype (p < 0.0001) and age (p < 0.0001), and a significant age x genotype interaction (p < 0.0001). *N =* 5 for *Repo >* GFP and *Repo >* GFP + GFP(IR); *N =* 10 for all other groups. (B) An equivalent analysis demonstrates that hTDP-43 has a similar effect in the neuronal cells of the *Drosophila* mushroom body using the *OK107-Gal4* driver, but with a later age of onset than hTDP-43 expression in glial cells. Quantification of GFP signal for each group is shown in the appropriate bar graph as in (A). A two-way ANOVA reveals significant effects of genotype (p = 0.0054) and age (p < 0.0001), as well as a significant age x genotype interaction (p = 0.0021). *N =* 5 for *OK107* > GFP and *OK107* > GFP + GFP(IR); *N =* 10 for all other groups. (C) Co-expression of hTDP-43, but not GFP, in the photoreceptor neurons of the fly eye under the *GMR-Gal4* driver interrupts the ability of a Dcr-2 processed IR to silence the endogenous *white+* pigment gene with an age of onset similar to that observed with neuronal expression of hTDP-43 in the CNS under *OK107-Gal4,* resulting in characteristic clusters of red-pigmented ommatidia. *N =* 5 for *GMR >* w(IR) + Gal80^ts^ OFF and *GMR >* w(IR) + Gal80^ts^ *ON; N = 20* for all other groups.

Although we have yet to identify which step of the siRNA pathway is disrupted by hTDP-43 expression, it is not simply due to loss of expression of *Dcr-2* or *Ago2,* the two major effectors of siRNA-mediated silencing in *Drosophila* (89-92). qPCR of whole head tissue demonstrated that hTDP-43 expression in both neurons and glial cells does not affect absolute expression levels *of Dcr-2* (Fig S7C.1) *or Ago2* (Fig S7C.2) at either 2-4 or 8-10 days post-eclosion, therefore down-regulation of these molecules is not responsible for the observed de-suppression *of gypsy.* In fact, in the case of genetic controls and flies expressing hTDP-43 in neurons, *Dcr-2* and *Ago2* levels actually increase with age beginning at 21-23 days post-eclosion and persisting into old age (40-42 days old), suggesting that down-regulation *of Dcr-2* and *Ago2* likewise cannot explain the later elevation *of gypsy* expression observed in these genotypes (Fig S7D.1-S7D.3). On the other hand, small-RNA seq from head tissue of flies expressing hTDP-43 under the glial *Repo-Gal4* driver reveals a relative reduction specifically in levels of antisense siRNAs among the subset that target RTEs whose expression is elevated in the RNAseq data (Table S5; Fig. S8A,B). This is suggestive of a defect in either biogenesis or stability of the siRNAs that target these RTEs. We favor a model (Fig 7) in which TDP-43 protein pathology interferes with siRNA biogenesis and/or function, resulting in deterioration of siRNA-mediated silencing accompanied by activation of RTE expression. The resulting increase in RTE expression may lead to accumulation of DNA damage resulting from RTE activity induced by TDP-43 pathology, in turn activating Loki/Chk2 signaling and leading to programmed cell death (Fig. 7).

**Fig 7.**
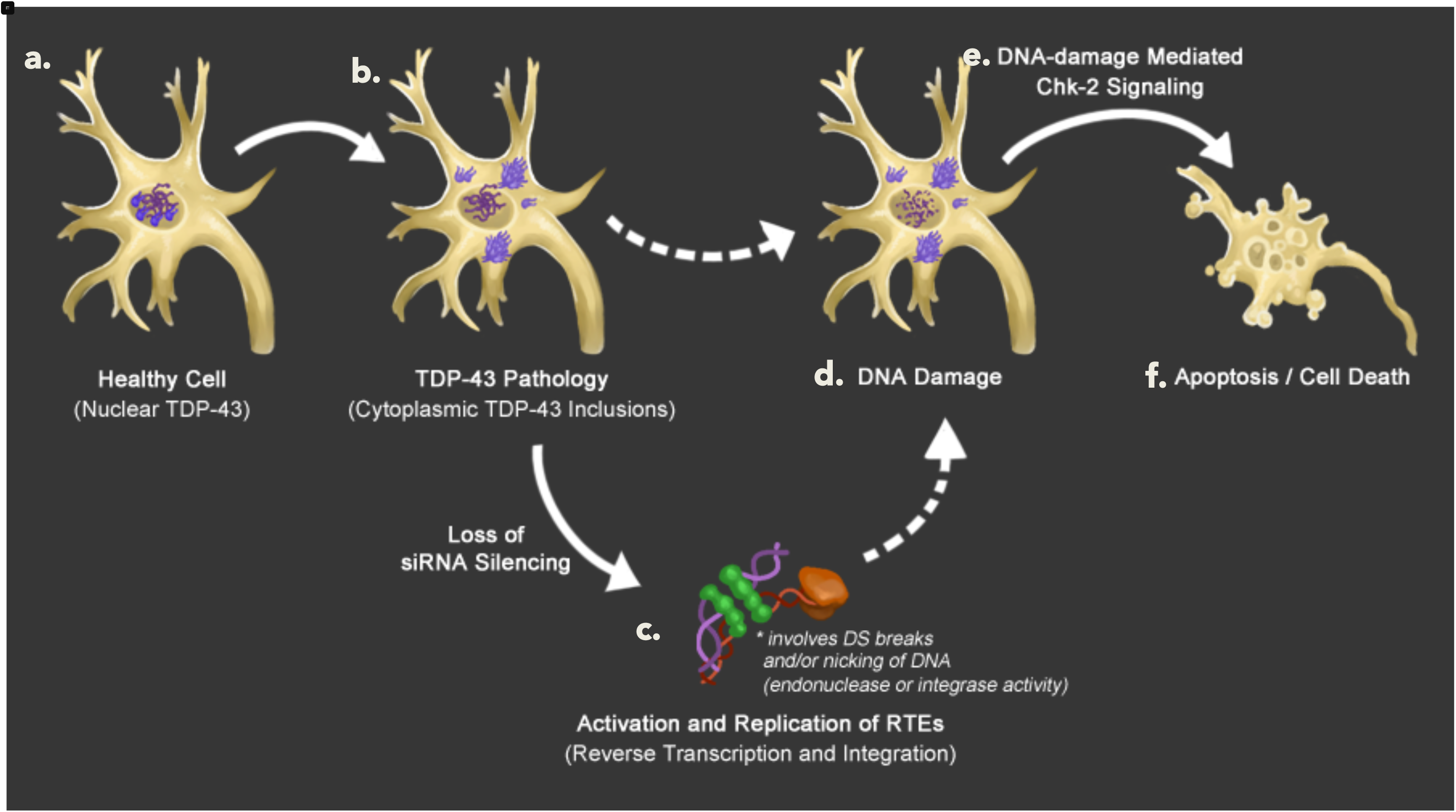
A cellular model for RTE mediated DNA damage and apoptotic cell death in TDP-43 pathology. In a healthy cell TDP-43 protein is predominantly found in the nucleus with the capacity to shuttle between the nucleus and the cytoplasm. This pleiotropic protein plays many important roles in normal RNA metabolism in both cellular compartments (A). Cells that experience TDP-43 pathology exhibit accumulation of TDP-43 protein in dense cytoplasmic inclusions and clearance from the nucleus (B). This is accompanied by rapid deterioration of siRNA-mediated silencing, as well as activation of RTE expression (C). The apoptotic cell death (F) induced by TDP-43 pathology is largely mediated by Loki/Chk-2 signaling (E). In the case of hTDP-43 pathology in fly glial cells in the CNS, expression of *the gypsy* RTE contributes to hTDP-43-induced cell death. Based on this observation, as well as the known cell biological role of Loki/Chk-2, we infer that the apoptosis induced by TDP-43 pathology is largely the result of Chk-2 activation in response to TDP-43-induced DNA damage (D), and that this DNA damage is at least partially incurred by TDP-43’s effects on RTEs (C).

## DISCUSSION

We previously reported bioinformatic predictions of a physical link between TDP-43 protein and RTE RNAs in rodent and in human cortical tissue (52). Here we provide *in vivo* functional evidence in *Drosophila* that TDP-43 pathological toxicity is the result of RTE activity generally and, in glial cells, expression of the *gypsy* ERV specifically. This finding is parsimonious with reports of high levels of reverse transcriptase activity in serum and CSF of HIV-negative ALS patients and their blood relatives (57-59), and of accumulation of transcripts and protein of HERV-K, a human ERV of the *gypsy* family, in the CNS of ALS patients (48, 53). It also is notable that accumulation of virus-like inclusions have been detected by electron microscopy in both neurons and glia of the frontal cortex of one ALS patient with extended prolongation of life via artificial lung ventilation (98). Furthermore, our findings are complementary to those documenting progressive motor dysfunction in transgenic mice expressing HERV-K ENV protein, one of the three major open reading frames of this human RTE (53). However, the findings reported here provide the first demonstration that an endogenous RTE causally contributes to physiological deterioration and cell death in TDP-43 protein pathology. Additionally, our findings indicate that reverse transcriptase enzymatic activity contributes to the toxicity of TDP-43 induced RTE expression, and that toxicity is largely mediated by DNA damage-induced cell death. Finally, we demonstrate that TDP-43 pathology leads to erosion of the post-transcriptional gene silencing mechanisms that are broadly responsible for RTE repression, which is accompanied by elevated expression of a panel of RTEs. These findings are in agreement with our previous observations that TDP-43 protein normally exhibits widespread interactions with RTE transcripts in rodent and human cortical tissue and that these interactions are selectively lost in cortical tissue of FTLD patients (52), as well as a report that knocking out the *C. elegans* ortholog of hTDP-43 results in broad accumulation of transposon-derived RNA transcripts and double stranded RNA (99).

TDP-43 has frequently been reported to co-localize with the major siRNA pathway components, DICER and Argonaute, in both cell culture and human patient tissue. Indeed such co-localization is commonly detected in stress granules (SGs), cytoplasmic foci for modulating mRNA translation that materialize in response to cellular stress (32, 45, 100-103). SGs are observed in pathological ALS and FTLD patient tissue, and can be induced in neuronal cell culture via overexpression of mutant and wild-type hTDP-43 as well as two other ALS linked genes, SOD1 and FUS, suggesting that they may represent a common downstream mechanism of pathological progression (45). Recent work by Emde et al. (2015) demonstrates that SG formation both in response to cellular stressors, or overexpression of ALS-linked genes including hTDP-43, inhibits DICER processing of pre-miRNAs to mature miRNAs. This signature is detectable in both sporadic and familial ALS spinal column motor neurons as a dramatic global reduction in mature miRNAs in comparison to control tissue (45). These findings are in accordance with previous work by Kawahara and Mieda-Sato (2012), which showed that loss of hTDP-43 function itself inhibits cytoplasmic miRNA processing by DICER for at least a subset of miRNAs (43). In mammals, the same DICER and Argonaute proteins process both miRNAs and siRNAs (104). Therefore, the effects of SG formation and hTDP-43 manipulation on DICER function may affect siRNAs just as dramatically as miRNAs, however the effects of TDP-43 expression on siRNA function in mammals have as yet to be investigated. In contrast with the mammalian system, siRNAs and miRNAs in *Drosophila* are processed largely via distinct pathways – Dcr-1/Ago1 and Dcr-2/Ago2, respectively (104). This disparity provided an opportunity for us to engineer an *in vivo* sensor to investigate the effects of TDP-43 on the siRNA system separate from its effects on miRNA biogenesis. While production of some miRNAs is disrupted in both animal models of ALS and human patient tissue, our data clearly demonstrate that in *Drosophila,* pathological TDP-43 expression disrupts the siRNA function of the DICER/Ago pathway, which then drives cellular toxicity, dramatic physiological deterioration, and premature death via loss of control of RTEs.

Unregulated RTE expression is known to be highly toxic in other biological contexts for a number of reasons, including accumulation of toxic RNAs, creation of harmful mutations, and accumulation of DNA damage. In the case of *gypsy* we demonstrate that this RTE is capable of actively mobilizing in response to TDP-43 pathology, and that it has a causal impact on cell death and physiological decline in the animal’s health. We also identify DNA damage-induced cell death, mediated by Chk2 activity, as a major contributing factor in the toxicity of TDP-43 both at the cellular and organismal level. Importantly, *gypsy* is not the only RTE whose expression we found to be increased in response to hTDP-43 expression. We in fact observe a panel of RTEs that exhibit elevated expression, with some variation in this profile when hTDP-43 is expressed solely in neurons or glia. This is consistent with the observation that knocking down *gypsy* expression only partially suppresses the toxicity of TDP-43, whereas blocking *loki (chk2)* expression leads to a near complete suppression of the effects of hTDP-43 expression on cell death and lifespan reduction. The involvement of DNA damage-induced cell death suggest that *gypsy* and likely other RTEs may be successfully or abortively inserting into genomic DNA, although we are mindful of the fact that increased levels of RTE proteins and RNAs may themselves be cytotoxic, as is observed with *the Alu* RTE in macular degeneration (50).

In the case of HERV-K, it has recently been shown (53) that TDP-43 binds directly to the LTR at the DNA level, thereby activating transcription of HERV-K. Our results establish, however, that TDP-43 pathology also compromises the siRNA-mediated gene silencing system, which is the major post-transcriptional genomic defense against RTEs in somatic tissues. The mechanisms by which TDP-43 protein pathology disrupts siRNA silencing remain to be investigated, but we favor the idea that it involves direct interactions between TDP-43 and the siRNA protein machinery (43, 45), and our previous findings also suggest direct interaction with RTE RNAs (52). The disruptive impact of TDP-43 on the siRNA system points to a general loss of RTE silencing – as opposed to activation of a specific element such as the *gypsy* ERV (or HERV-K) – as the major contributing factor in hTDP-43-related neurophysiological deterioration. This conclusion is supported by our RNA-seq data which shows a broad and general increase in RTE expression in head tissue of *Drosophila* expressing either neuronal or glial hTDP-43, as well as a pronounced reduction specifically in antisense siRNAs which target the RTEs we observe to exhibit increased expression in response to hTDP-43 expression. In accordance with this notion, we have previously shown in *Drosophila* that mutation of Ago2, a major effector protein of the siRNA system, results in activation of several different RTEs in brain tissue and causes rapid age-related cognitive decline and shortened lifespan (19).

Like *Drosophila,* the human genome contains more than one type of functional RTE. In addition to HERV-K, the human genome contains on the order of 100 fully active copies of L1 RTEs, and a far higher number of non-autonomous elements that replicate in *trans* by capitalizing on the protein machinery encoded by L1s (105). Moreover, abnormally high levels of expression of several different RTE families has been reported across a suite of neurodegenerative diseases (48-55), and there is accumulating evidence suggesting RTEs generally become active with advanced age in a variety of organisms and tissues (13-18), including the brain (19). Previous studies make the case that this effect may result from age-related loss of transcription-level heterochromatic silencing (16, 18). Our finding that TDP-43 erodes post-transcriptional, siRNA-mediated RTE silencing therefore raises an intriguing hypothesis regarding the synergy between age and TDP-43 pathology on RTE activation, particularly when the reinforcing action of siRNAs on heterochromatin is taken into consideration (1, 18). This potential synergy, in conjunction with the replicative capacity encoded by RTEs, leads us to posit the “retrotransposon storm” hypothesis of neurodegeneration. We envision that loss of control of RTE expression and replication leads to a feed-forward mechanism in which massive levels of activity drive toxicity and degeneration in the nervous system. Our findings are not in conflict with a wealth of data that have implicated effects of TDP-43 pathology on splicing, RNA stability, translation, and miRNA biogenesis (26, 27, 36, 37, 43-45)., and it will be important to conceptually integrate our findings with these other aspects of TDP-43 pathology. But the direct impact we observe on cell death highlights the importance of investigating the contribution of siRNA dysfunction and RTE toxicity in TDP-43-mediated pathogenesis, and may indicate a promising common avenue for novel therapeutic targets in both familial and sporadic cases of ALS.

## MATERIALS AND METHODS

### Fly Stocks

All transgenic fly stocks used, with the exception of w(IR) and *GMR-Gal4,* were backcrossed into our in-house wild type strain, the Canton-S derivative *w*^*1118*^ (*isoCJ1*) (106), for at least five generations to homogenize genetic background. The GFP, *OK107-, ELAV-,* and *Repo-Gal4* lines (107), as well as the hTDP-43 (66) and *gypsy*(IR) (55) lines, are as reported previously. The *GMR-Gal4*, *Gal80*^*ts*^, w(IR), GFP(IR), and tdTomato lines were acquired from the Bloomington Drosophila Stock Center (stock numbers: 43675, 7019, 25785, 9331, and 32221; respectively), and the loki(IR) line was acquired from the Vienna Drosophila Resource Center (108) (stock number: v44980). Flies were cultured on standard fly food at room temperature unless otherwise noted.

### Bleach Treatment of Embryos

All fly stocks used for lifespan analysis and longitudinal qPCR experiments were double dechorionated by bleach treatment in order to remove exogenous viral infection (19). Briefly, 4-hour embryos were collected and treated with 100% bleach for 30 min to remove the chorion. Treated embryos were washed and subsequently transferred to a virus-free room equipped with ultraviolet lights to maintain sterility. This was repeated for at least two successive generations and expanded fly stocks were tested via qPCR of whole flies to ensure Drosophila C Virus (DCV) levels were below a threshold of 32 cycles.

### RNA-Seq and small RNAseq Library Preparation

Fly heads were collected for each genotype and total RNA was purified with Trizol (Invitrogen). RNA-Seq libraries were constructed using the NuGEN Ovation Drosophila RNA-Seq kit including DNase treatmen with HL-dsDNase (ArcticZymes Cat. # 70800-201) and cDNA fragmenation using the Covaris E220 system according to manufacturer specifications. After amplification, library quality was measured using the Agilent Bioanalyzer system and quantity was determined using Life Technologies Qubit dsDNA HS Assay kit (for use with the Qubit 2.0 Fluorometer). Prior to sequencing, pooled libraries were quantified using the Illumina Library Quantification kit (with Universal qPCR mix) from Kapa Biosystems and a 1.2 pM loading concentration was used for PE101 on the Illumina NextSeq500 platform. Small RNAs were cloned using TruSeq (Illumina) approach with modifications described in Rozhkov 2015 (109). Briefly, all small RNA libraries were constructed from 15-20 ug of TRIzol isolated total RNA. 18-29 nucleotide long small RNAs were size selected on 15% PAGE-UREA gel. After 3’-and 5’-adapter ligation, subsequent gel purification steps and reverse transcription cDNAs were PCR amplified with barcoded primers. PCR products were size selected on 6% PAGE gel, and quantified on Bioanalyzer. Loading concentrations were determined using the NEBNext Library Quant kit for Illumina (NEB). The libraries were sequenced on NEXTSeq platform.

### Sequencing, Mapping and Annotation

RNA-seq libraries were run on an Illumina NextSeq (paired end 101). Reads were mapped to the *Drosophila* dm3 genome with STAR (110) allowing up to 4 mismatches and a maximum of 100 multiple alignments. To estimate the pileup along *gypsy* element, reads were mapped to the *gypsy* consensus sequence (GenBank accession: M12927) using Bowtie (111) with up to 2 mismatches. Reads were annotated based on genomic locations against ribosomal RNAs, transposable elements (FlyBase), and RefSeq genes (UCSC genome database RefSeq track).

### Transcriptome analysis

Reads mapped to ribosomal RNAs were removed from each library. For the remaining reads, expression abundance estimation and differential expression analysis were performed using the TEtranscripts package (74). Reads for each library were normalized based on library size, e.g., reads per million mapped (RPM). Statistically significant differences were taken as those genes/transposons (TEs) with a Benjamini-Hochberg corrected P-value < 0.05, as calculated by DESeq (112). Biological replicates were averaged for the purpose of estimating pileup along consensus TEs.

### Enrichment analysis for KEGG ALS pathway

The ortholog pairs of fly and human were predicted with the Drosophila Integrated Ortholog Prediction Tool (DIOPT), which integrates the ortholog predictions from 11 existing tools (113). The pairs with the “high” rank, defining as the best match with both forward and reverse searches and the DIOPT score is at least 2, were selected. Then we performed Fisher’s exact test to check if the differential expression fly genes are enriched in the KEGG ALS human genes with identifiable fly orthologs.

### Lifespans

Male flies were used for all lifespan assays since the majority of glial-expressing hTDP-43 flies that escape their pupal cases are male. Flies were housed 15 to a vial with a total of 75 flies per genotype and flipped into fresh food vials every other day. All vials were kept on their side in racks for the duration of the experiment. Lifespan experiments were performed blind.

### NRTI Food Preparation and Lifespans

Tenofovir disoproxil fumarate (TDF; Selleck Chemicals, CAS 147127-20-6), Zidovudine (AZT; Sigma-Aldrich, CAS 30516-87-1), and Stavudine (d4T; Sigma-Aldrich, CAS 3056-17-5) were prepped as solutions using dimethyl sulfide (DMSO) as solvent. Standard fly food was melted and cooled to a liquid, and NRTI solutions were added just before solidification to give final concentrations of 0 (vehicle alone control), 1, 5, 10, and 15 μM of each NRTI in a total volume of 0.2% DMSO, then stored at 4 °C for a maximum of 2 weeks until use. Lifespan was monitored in all male flies with 20 flies per vial for a total of 100 flies monitored per genotype per treatment. Lifespans were performed as above.

### CAFE Assays

To quantify and determine the consistency of feeding across treatments, Capillary Feeder (CAFE) Assays (114) were performed. Each replicate consisted of five male wild-type flies contained in a 1.5 mL microfuge tube chamber with 1% agarose in bottom to maintain humidity and two 5 μL disposable calibrated pipets (VWR, 53432-706) providing the media solution (5% sucrose / 5% autolyzed yeast extract, Sigma-Aldrich), inserted though the tube cap. Flies were acclimated with untreated solution *ad libitum* for 24 hours. Measurements were initiated one hour following the switch to experimental solutions and monitored for a total duration of 24 hours. Experimentally treated solutions were prepared to match the concentrations of each NRTI treatment used in solid food for lifespan analysis, as well as an additional solution of high concentration for each NRTI (100 μM) and a solution of vehicle alone control (0.2% DMSO). Displacement due to evaporation was controlled for by subtracting measurements from fly-less CAFE chambers with the vehicle solution set up in parallel to the experimental assays. CAFE assays with untreated solution were also set up in parallel and controlled for evaporation.

### Locomotion Behavior Assay

Locomotion behavior was assayed using the classic Benzer counter current apparatus as in Benzer, S., 1967 (115), with the following modifications: freshly eclosed flies were transferred into glass bottles with food and a paper substrate and plugged with foam stoppers. Flies were transferred to fresh bottles every 48 hours until they reached the appropriate age for locomotion assays. The Benzer assay was conducted in a horizontal position with a fluorescent light source to measure phototaxis. Locomotion assays were performed blind.

### qPCR and TaqMan Probes

Tissue preparation, cDNA synthesis and qPCR were performed as previously described (19) using the Applied Biosystems StepOnePlus Real-Time PCR System. Heads of 75-100 flies were used for each biological replicate unless otherwise noted. All TaqMan Gene Expression Assays were acquired from Applied Biosystems and used the FAM Reporter and MGB Quencher. The inventoried assays used were: *Act5C* (assay ID Dm02361909_s1), *Dcr-2* (assay ID Dm01821537_g1),Ago2 (assay ID Dm01805433_g1), *TARDBP* (assay ID Hs00606522_m1), *TBPH* (assay ID Dm01820179_g1), and *loki* (assay ID Dm01811114_g1). All custom TaqMan probes were designed following the vendor’s custom assay design service manual and have the following assay IDs and probe sequences: *gypsy ORF2* (assay ID AI5106V; probe: 5’-AAGCATTTGTGTTTGATTTC-3’), *gypsy ORF3* (assay ID AID1UHW; probe: 5’-CTCTAGGATAGGCAATTAA-3’), and *DCV* (assay ID AIPAC3F; probe: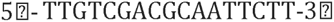).

### Whole Mount Immunohistochemistry and GFP Imaging

Dissection, fixation, immunolabelling, and confocal imaging acquisition were executed as previously described (116). The ENV primary antibody was used as described in Li, *et al.* 2013 (19, 76). For TDP-43 immunohistochemistry, the primary full length human TDP-43 antibody (Protein Tech, 10782-2-AP) was used at a 1:100 dilution, and the primary pSer409 phosphorylated human TDP-43 antibody (Sigma Aldrich, SAB4200223) was used at a 1:500 dilution separately in conjunction with a 1:200 dilution of an Alexa Fluor 488-conjugated secondary antibody (Thermo Fisher Scientific, A-11070). Repo co-labeling was performed using a 1:200 dilution of primary antibody (Developmental Studies Hybridoma Bank, 8D12) and a 1:200 dilution of a Cy3-conjugated secondary antibody (Molecular Probes, A10521). DAPI co-staining was performed after a brief wash in 1x PBS immediately subsequent to secondary antibody staining using DAPI Dilactate (Thermo Fisher Scientific, D3571) as per manufacturer specifications. All brains co-stained with DAPI were imaged on a Zeiss LSM 780 confocal microscope using a UV laser and the Zeiss ZEN microscope software package.

### GFP Quantification

The gain on the confocal microscope was set using the positive control *(Repo >* GFP or *OK107 >* GFP) and kept consistent across all subsequent brains imaged. The GFP signal of the median 10 optical sections of the appropriate structures (either the full brain for *Repo* or both lobes of the calyx for *OK107,* respectively) was calculated using ImageJ software, as previously described (117). These ten values were then averaged, and this number used as a representation for each individual brain. 5-10 brains were analyzed per group.

### TUNEL-positive Nuclei Detection and Quantification

For TUNEL staining, the *In Situ* Cell Death Detection Kit, TMR red (Roche, 12156792910) was used. The same dissection, fixation, and penetration and blocking protocol used for antibody staining was followed (116), at which point the brains were transferred to the reaction mix from the kit for 2 hours at 4 °C followed by 1 hour at 37 °C. Brains were then washed, mounted, and imaged as previously described (116). For imaging, the gain on the confocal microscope was set using the positive control (*Repo* > hTDP-43) and kept consistent across all subsequent brains imaged. A projection image was generated using the middle 50 optical slices from the z-stack image of the whole brain. This projection image was then thresholded using the maximum entropy technique (See: (118)) via the Fiji plug-in for ImageJ software, and the subsequent binary image was subjected to puncta quantification using ImageJ software. Puncta quantification was thresholded for puncta greater than 3 pixels to reduce the likelihood of counting background signal. The total number of puncta counted was then used as a representation for the number of TUNEL-positive nuclei for each brain in subsequent statistical analysis. 7-12 brains were analyzed per group. Quantification *of gypsy* ENV immunoreactive puncta was performed in the same manner.

### *Drosophila* Eye Imaging

Flies of the appropriate age and genotype were placed at -70 °C for 25 minutes and then kept on ice until immediately prior to imaging. Imaging was performed using a Nikon SMZ1500 stereoscopic microscope, Nikon DS-Vi1 camera and Nikon Digital Sight camera system, and the Nikon NIS-Elements BR3.2 64-bit imaging software package. The experiment was designed such that each group is balanced for the number of mini-white transgenes and heterozygous for genomic *white*^*+*^.

### Tranmission Electron Microscopy

*Drosophila* heads were removed, the cuticle removed and the brains fixed overnight in 2% paraformaldehyde and 2% glutaraldehyde in 0.1 mol/L PBS. Samples were rinsed in distilled water and post-fixed for one hour in 1% osmium tetroxide in 1.5% potassium ferrocyanide in distilled water. Next, the samples were dehydrated in a graded series of ethanol and the final 100% ethanol was replaced with a solution of absolute dry acetone (Electron Microscopy Sciences, Hatfield PA). The samples were then infiltrated with agitation for one hour in an equal mixture of acetone and Epon-Araldite resin, followed by infiltration with agitation overnight in 100% resin. Samples were transferred to embedding capsules with the posterior head facing towards the bottom of the capsule and the resin was polymerized overnight in a vented 60 °C oven. Thin sections were made from the mushroom body region and collected on Butvar coated 2mm x 1mm slot grids (EMS) and the sections were counterstained with lead citrate. Thin sections were imaged with a Hitachi H700 transmission electron microscope and recorded on Kodak 4480 negatives that were scanned with an Epson V750 Pro Scanner at 2400 DPI. 3 individual brains processed for *Repo* / +, 4 individual brains processed for *Repo >* hTDP-43; many images collected of each brain.

### Statistics

For qPCR data, the p-values of all data sets with only two groups were calculated using an unpaired t-test. Where an effect of age for more than two time points within one genotype was determined, a one-way ANOVA was performed, and where multiple ages and genotypes are represented a two-way ANOVA was performed; the results are reported in the figure legends. All pairwise comparisons for qPCR reported in the figures were corrected using the Bonferroni method for multiple comparisons. For both the locomotion data and the GFP quantification, p-values were reported using the Sheffé method; ANOVA results are reported in the figure legends. The CAFE assay is similarly reported, with pairwise comparisons between treatments being made using one-way ANOVA with Tukey^’^s Multiple Comparisons test. Survival analyses for the lifespan curves were performed using the Kaplan-Meier method, and the Gehan-Breslow-Wilcoxon test were used to compare survival curves. All pairwise comparisons for lifespan curves were corrected using the Bonferroni method. Sample sizes were selected based on standard practices in the literature. No randomization was employed in this study.

## ACKNOWLEDGEMENTS

We would like to thank Jef Boeke for the 7B3 hybridoma cell line and Carmelita Bautista at the CSHL shared resources for ascites production. We thank Magalie Lecourtois for the hTDP-43 fly line and Peng Jin for the *gypsy(IR)* line. Stocks obtained from the Bloomington Drosophila Stock Center (NIH P40OD018537) and the Vienna Drosophila Resource Center (108) were also used in this study. We would also like to thank Will Donovan for aid in the whole mount TUNEL staining assay, as well as Rob Martienssen, Steve Shea, Meng-Fu Shih and Yung-Heng Chang for helpful discussion.

### AUTHOR CONTRIBUTIONS

JD and LK conceived and designed most of the experiments and performed most of the analyses. LK performed most of the experiments, with the following exceptions: NC performed the locomotion behavioral assays and the majority of the ENV immunostaining. RBM performed the TDP-43 antibody immunostaining. SH performed tissue preparation and imaging for TEM. KM performed the NRTI food preparation and lifespan analysis. DT performed the qPCR to detect *loki* and *gypsy* levels in the RNAi experiments. LP prepared the RNAseq libraries with help from NR and WWL conducted the RNAseq analysis under the guidance of MH. NR performed the small-RNAseq library preparation and WWL conducted the sRNAseq analysis under guidance of MH. YHC and LP constructed the gypsy-G2M reporter, YHC performed the in vivo imaging of the G2M reporter and RK performed the S2 cell imaging. This manuscript was written by JD and LK with comments from the other authors.

## SUPPORTING INFORMATION

Supplemental information including 5 Supporting Figures and 5 Supporting Tables can be found with this article online at:

S1 Fig. Differentially expressed genes in Repo-TDP-43 heads enriches for fly orthologs of ALS-KEGG gene set. Among the KEGG gene set annotated as functionally related to ALS pathways. Although this is by no means a complete list of genes that have been implicated in ALS in the literature, it represents a set of gene pathways with known involvement in ALS. 20 of the ALS KEGG gene set contain clear Drosophila orthologs and a significant fraction of these (11/20) are identified as differentially expressed in our RNAseq from *Repo>TDP-43* heads.

**S2. Fig.*gypsy* expression turns on stochastically in young brains and reaches peak expression in the population at mid-adulthood.**

(A) Transcript levels of *gypsy ORF3 (Env)* as detected by qPCR on whole head tissue of flies expressing (A.1) neuronal hTDP-43 *(ELAV>* hTDP-43), or genetic controls: (A.2) *ELAV/* + and (A.3) hTDP-43 / +. *gypsy ORF3* transcript levels display an increase by 21-23 days post-eclosion that drops back down by 40-42 days post-eclosion, regardless of genotype. In all cases transcript levels have been normalized to *Actin,* and the aged cohort (8-10 days; 21-23 days; 40-42 days) are represented as a fold change over an appropriate young (2-4 day) cohort that has been processed in parallel. Unpaired t-tests have been used to calculate p-values for each aged cohort with its matched young cohort, while p-values comparing aged cohorts within genotypes have been calculated using the Bonferroni method for multiple comparisons. For all three genotypes a one-way ANOVA shows a significant effect of age *on gypsy ORF3* transcript levels between the aged cohorts (*ELAV* > hTDP-43, p < 0.0001; *ELAV /* +, p < 0.0001; hTDP-43 / +, p = 0.0346). *N =* 5 for all groups. (A.4). qPCR of whole head tissue reveals that the presence of the hTDP-43 transgene alone with no Gal4 driver results in elevation of *gypsy* ORF2 transcript levels. N = 6 for both groups. Quantitative genomic PCR (A.5) reveals that the wild type, Elav-Gal4 and Repo-Gal4 lines have comparable levels of *gypsy* DNA copy number, and (A.6) mRNA expression levels. The *UAS-hTDP-43* parental line exhibits marginally higher levels of *gypsy* genomic DNA (A.5), although this cannot explain the difference between expression in *ELAV>* hTDP-43 vs *Repo>* hTDP-43. (B) Quantification of immunoreactive puncta averaged across the central 10 optical slices of brains of flies expressing hTDP-43 in glial cells *(Repo >* hTDP-43) and genetic controls *(Repo* / + and hTDP-43 / +) aged to 10 days and whole-mount immunostained using the *gypsy* ENV monoclonal antibody in a separate experiment from Fig 1C. A one-way ANOVA reveals that there is no difference in ENV immunoreactivity between the two genetic control groups *(Repo /* + and hTDP-43 / +; p = 1.0000), but that 10 day old flies that express hTDP-43 in glial cells *(Repo >* hTDP-43) display significantly more ENV immunoreactive puncta than flies carrying the UAS-hTDP-43 transgene (hTDP-43 / + ; p < 0.0001) or Repo-Gal4 alone *(Repo /* + ; p < 0.0001). Displayed as means + SEM; *N =* 8 for all groups. (C) Projections through whole-mount brains immunolabeled with *agypsy* ENV monoclonal antibody demonstrate that *gypsy expression* turns on post-developmentally, with very little *gypsy* expression immediately following eclosion (0 Days) in genetic controls (hTDP-43 / + and *Repo /* +) and in flies expressing hTDP-43 in glia *(Repo >* hTDP-43). Expression turns on stochastically at 3 days post-eclosion only in the CNS of flies expressing glial hTDP-43. Replicates of 3-day old *Repo >* hTDP-43 brains illustrate the variability of *gypsy* expression at this early time point. *N =* 2 for all 0 Day groups; hTDP-43 / +, 3 Day, *N =* 3; *Repo /* +, 3 Day, *N =* 4; *Repo >* hTDP-43, 3 Day, *N =* 7.

S3. Fig. A reporter of gypsy mobilization.

(S3A) Design of the Gypsy-G2M reporter takes advantage of the replication mechanism of retroviruses. Because of two separate template switches, sequences from the right half of the 5’LTR are used to complete replication of the 3’LTR. And likewise, sequences from the left half of the 3’LTR are used as template to complete the 5’LTR. We placed a UAS element that lacked a reporter into the 5’LTR and a reporter lacking a promoter (myrGFP-P2A-H2BmCherry, G2M) into the 3’LTR. After replication and re-insertion into the genome, the UAS element is placed upstream of the G2M reporter. G2M encodes a membrane GFP, followed by the P2A self cleaving peptide, followed by a nuclear-mCherry. Thus expression of G2M labels the nucleus with mCherry and the membrane with GFP, as shown in Drosophila S2 cells. Details of this reporter design will be described elsewhere. (S3B) Replication of gypsy-G2M was monitored in transgenic flies by detection of mCherry nuclear label under control of the Repo-Gal4 line. Expression of mCherry (red) was seen in glial cells that were co-labeled with an antibody against the repo protein (Blue), which marks glial nuclei. This reporter system reveals a significant (S3B,C) age dependent increase in gypsy replication in control flies that carry only a UAS-lacZ cassette (S3B left panels compare day 2 age with day 5-7 age, S3C for quantification). This age-related expression is consistent with our prior observations of gypsy replication during aging using a different reporter (19).Co-expression of hTDP-43 under Repo-Gal4 causes a significant increase in mCherry labeling at both ages (S3B right panels; S3C for quantification). In addition, the accumulation of mCherry labeled nuclei also depends upon gypsy expression because co-expression of the gypsy-IR significantly reduces the reporter expression (S3C).

**S4 Fig. Characterizing hTDP-43 expression.**

(A) TUNEL staining reveals very little apoptotic activity when hTDP-43 is expressed in the mushroom body under *OK107-Gal4,* even when the animals are aged to 30 days post-eclosion. Mushroom bodies marked by co-expression of GFP, shown in green; TUNEL staining shown in red. *OK107* > GFP, 5 Day, *N =* 3; *OK107* > GFP + hTDP-43, 5 Day, *N =* 5; *OK107* > GFP, 30 Day, *N =* 5; *OK107 >* GFP + hTDP-43, 30 Day, *N =* 4. (B) Full length human TDP-43 (green) can be detected by immunolabelling in the brains of flies expressing glial hTDP-43 under the *Repo-Gal4* driver at 21-23 days post-eclosion, and co-localizes with Repo (red) immunoreactivity (left). *Repo / + (N =* 4); *Repo >* hTDP-43 (*N =* 4). Immunoreactivity for a disease-specific phosphorylated isoform of the protein (pSer409) can also be readily detected and co-localizes with Repo (right). *Repo /* + (*N =* 7); *Repo >* hTDP-43 (*N =* 4). A 63x blow-up is shown in the pop-out. (C) Both the full-length (left) and disease specific (right) isoforms of hTDP-43 (green) are mainly observed in the cytoplasm and vacate the nucleus (visualized by DAPI co-staining, shown in blue). Arrowheads indicate the hTDP-43-filled cytoplasm of a cortical glial cell wrapped around several neuronal nuclei in the neuropil of flies expressing glial hTDP-43. For full length hTDP-43 antibody, *Repo / + (N=* 6), *Repo >* hTDP-43 (*N =* 13); for pSer409 phosphorylated hTDP-43 antibody, *Repo / + (N=* 4), *Repo >* hTDP-43 (*N =* 9). (D) qPCR of whole head tissue demonstrates that transcript levels of *hTDP-43* diminishes under *Repo-Gal4* from 2-4 days to 8-10 days. Transcript levels normalized to *Actin* and displayed as fold change relative to 2-4 day old flies (means + SEM). *N =* 6 for all groups. (E) A similar effect of age on hTDP-43 expression is observed in neurons under *ELAV-Gal4,* and continues to drop off by 40-42 days post-eclosion. A one-way ANOVA shows a significant effect of age (p < 0.0001). *N =* 6 for all groups.(F) qPCR of whole head tissue demonstrates that expression of hTDP-43 does not effect levels of the Fly ortholog, *TBPH,* regardless of cell type of expression. Transcript levels normalized to *Actin. N =* 4 for the hTDP-43 / + group, *N =* 5 for all other groups.

**S5 Fig. NRTI administration does not affect fly feeding behavior and expression of IR constructs individually does not affect lifespan or hTDP-43 expression. However, NRTI administration extends lifespan in flies expressing hTDP-43 in glia.**

(A) qPCR on head tissue demonstrates that expressing an IR directed against *gypsy ORF2 (gypsy(IR))* in neurons *(ELAV> gypsy(IR))* or glia *(Repo > gypsy(IR))* effectively inhibits age-dependent elevation of *gypsy* transcript levels, and results in an ∼2.5-fold reduction at 28 days post-eclosion. *gypsy* transcript levels from head tissue of young (2-4 day) and aged (28 day) flies was normalized to *Actin* and displayed as fold change in 28 day old flies of each genotype relative to each respective young cohort (displayed as means + SEM). A one-way ANOVA shows a significant effect of genotype (p = 0.0182). *N =* 2-3 biological replicates generated from heads of 5 mL of flies for each group. (B) Expression *of gypsy(IR), loki(IR),* and GFP(IR) individually in glial cells under the *Repo-Gal4* driver does not significantly alter lifespan. (C) Expression *of gypsy(IR)* and *loki(IR)* individually in neurons under the *ELAV-Gal4* driver does not significantly alter lifespan. (D) The presence of each of the IR constructs alone without any Gal4 driver *(gypsy(IR) /* + ; *loki(IR) /* +; GFP(IR) / +) only moderately effects lifespan. (E) The capillary feeder (CAFE) assay demonstrates that neither the final concentration of vehicle (0.2% DMSO) alone, any of the final concentrations of D4T used in solid fly food in the lifespan analysis (Fig 4D; 1 ⎧M, 5 ⎧M, 10 ⎧M, or 15 ⎧M D4T in 0.2% DMSO), or an additional high concentration of D4T (100 ⎧M in 0.2% DMSO) significantly altered displacement by consumption by wild type flies of a liquid media solution (5% sucrose / 5% autolyzed yeast) (one-way ANOVA; p = 0.2137) over the course of 24 hours. Fly-less assay tubes containing vehicle solution alone were used to control for evaporation. Displayed as means + SEM; *N =* 6-8 for all D4T groups. (F) CAFE assay performed as in Fig S5E demonstrates that AZT does not significantly alter feeding behavior of wild type flies (p = 0.0595) at any of the concentrations used in solid fly food in lifespan analyses (Fig 4E; 1 ⎧M, 5 ⎧M, 10 ⎧M, or 15 ⎧M AZT in 0.2% DMSO) or at an additional high concentration of AZT (100 ⎧M in 0.2% DMSO). Displayed as means + SEM; *N =* 9-12 for all AZT groups. (G) Lifespan analysis shows that the NRTI Stavudine (D4T) partially suppresses the lifespan deficit induced by glial expression of hTDP-43 *(Repo >* hTDP-43) when supplied in solid fly food at final concentrations of either 5 ⎧M, p = 0.0450; or 10 ⎧M, p = 0.0186; in comparison to vehicle alone control (0 (M). (H) Lifespan analyses performed as in (D) for the NRTI Zidovudine (AZT). AZT partially suppresses the lifespan deficit exhibited by flies expressing hTDP-43 in glia *(Repo >* hTDP-43) when supplied at 5 (M (p = 0.0245) in comparison to a vehicle alone control (0 ⎧M).

**S6 Fig. Co-expression of IR constructs does not alter hTDP-43 expression level.**

(A) An equivalent analysis as described for (S5A) demonstrates that neuronal (*ELAV* > loki(IR)) and glial *(Repo > loki(IR))* expression of an IR directed against *loki (loki(IR))* effectively blocks the age-dependent elevation of *loki* transcript levels, resulting in an ∼2-fold reduction at 28 days post-eclosion. A one-way ANOVA shows a significant effect of genotype (p = 0.0039). *N =* 3-4 biological replicates. (B) Co-expression of GFP(IR) with hTDP-43 under Repo-Gal4 *(Repo >* hTDP-43 + GFP(IR)) does not significantly alter the number of TUNEL-positive nuclei detected compared to hTDP-43 expression alone under Repo-Gal4 *(Repo >* hTDP-43). *N =* 8 for *Repo >* hTDP-43 and *N =* 9 for *Repo >* hTDP-43 + GFP(IR); data normalized to *Repo >* hTDP-43. (C) qPCR for hTDP-43 expression *(TARDBP)* on whole head tissue demonstrates that co-expression of each of the IR constructs with hTDP-43 under Repo-Gal4 *(Repo >* hTDP-43 + GFP(IR), *Repo >* hTDP-43 + *gypsy(IR),* and *Repo >* hTDP-43 + *loki(IR),* respectively) does not significantly reduce hTDP-43 expression levels compared to hTDP-43 expression alone under Repo-Gal4 *(Repo >* hTDP-43) Fold change is displayed as the mean fold change relative to *Repo >* hTDP-43, while p-value represents the p-value of a two-tailed Student’s t-test in comparison to *Repo >* hTDP-43. *N =* 4 for all groups.

**S7 Fig. Turning on *white+* expression post-developmentally rescues red eye pigmentation in *Drosophila.* Loss of suppression of *gypsy* cannot be explained by hTDP-43- or age-dependent effects on siRNA effector molecules.**

(A) Schematic representation of the experimental design. (B) Representative images demonstrating that turning off w(IR) expression post-developmentally rescues red pigmentation of the Fly eye. *N =* 5 for all groups. (C) qPCR of whole head tissue demonstrates that reduced expression of (C.1) *Dcr-2* and (C.2) *Ago2* cannot account for loss of suppression *of gypsy* in flies expressing glial hTDP-43 *(Repo >* hTDP-43) at either 2-4 or 8-10 days post-eclosion. Transcript levels normalized to *Actin* and displayed as fold change relative to flies carrying the hTDP-43 transgene with no Gal4 driver (hTDP-43 / +) at 2-4 Days (means + SEM). For *Dcr-2,* a two-way ANOVA reveals an effect of age (p = 0.0006) but no effect of genotype (p = 0.1081); for *Ago2,* a two-way ANOVA also reveals an effect of age (p = 0.0258) but no effect of genotype (p = 0.1591). *N =* 8 for all groups. (D) qPCR of whole head tissue demonstrates that age-dependent changes in expression of *Dcr-2* (top) *and Ago2* (bottom) cannot account for age-dependent loss of suppression of *gypsy* in flies expressing neuronal hTDP-43 *(ELAV>* hTDP-43; D.1) or genetic controls: (D.2) *ELAV/* + and (D.3) hTDP-43 / +. All data analyzed as in (S1A.1-S1A.3); one-way ANOVA shows an effect of age across almost all groups *(ELAV>* hTDP-43, *Dcr-2,* p < *0.0001, Ago2,* p = 0.0269; *ELAV/ +, Dcr-2,* p < *0.0001, Ago2,* p = 0.0051; hTDP-43 / +, *Dcr-2,* p < *0.0001, Ago2,* p = 0.3967). *N* = 8 for all groups.

**S8. Fig. Expression of hTDP-43 results in a loss of siRNAs anti-sense to RTEs whose expression is increased. Sequencing of small-RNAs from heads detects siRNAs that are predicted to target broad range of RTEs. Among RTEs whose expression is elevated in Repo>TDP-43 (vs control), there is a selective decrease in anti-sense relative to sense stranded siRNAs. (A) scatter plots of log2Fold change in RNAseq vs log2Fold change in small-RNA seq. 3S18, mdg3 and gypsy exhibit elevated levels of expression in the RNAseq data (see Fig. 1), and also exhibit decreased anti-sense/sense siRNA ratio. This is in contrast to Burdock, whose RNA levels are unchanged and whose siRNA anti-sense/sense ratio is unchanged. (B) Overall, there is a significant decrease in anti-sense/sense ratio for the subset of siRNAs that map to TEs whose levels are altered by Repo>hTDP-43.**

**S1 Table. Mapping statistics for RNA-seq.**

**S2A Table. Abundance of gene-derived transcripts in head tissue of flies expressing pan-neuronal hTDP-43 compared to controls.**

**S2B Table. Abundance of transposon-derived transcripts in head tissue of flies expressing pan-neuronal hTDP-43**

**compared to controls.**

S2C Table: Fly orthologs of ALS KEGG gene set.

**S3A Table. Abundance of gene-derived transcripts in head tissue of flies expressing pan-glial hTDP-43 compared to controls.**

**S3B Table. Abundance of transposon-derived transcripts in head tissue of flies expressing pan-glial hTDP-43 compared to controls.**

**S4 Table: Expression levels (RPM) of fly TBPH and hTDP-43**

**S5 Table: Summary statistics for small RNA seq mapping**

## REFERENCES

1. Slotkin RK, Martienssen R. Transposable elements and the epigenetic regulation of the genome. Nature reviews Genetics. 2007;8(4):272–85.

2. Wallace NA, Belancio VP, Deininger PL. L1 mobile element expression causes multiple types of toxicity. Gene. 2008;419(1-2):75–81.

3. Garcia-Perez JL, Marchetto MC, Muotri AR, Coufal NG, Gage FH, O’Shea KS, et al. LINE-1 retrotransposition in human embryonic stem cells. Human molecular genetics. 2007;16(13):1569–77.

4. Kazazian HH, Jr. Mobile DNA transposition in somatic cells. BMC biology. 2011;9:62.

5. Baillie JK, Barnett MW, Upton KR, Gerhardt DJ, Richmond TA, De Sapio F, et al. Somatic retrotransposition alters the genetic landscape of the human brain. Nature. 2011;479(7374):534–7.

6. Coufal NG, Garcia-Perez JL, Peng GE, Yeo GW, Mu Y, Lovci MT, et al. L1 retrotransposition in human neural progenitor cells. Nature. 2009;460(7259):1127–31.

7. Evrony GD, Cai X, Lee E, Hills LB, Elhosary PC, Lehmann HS, et al. Single-neuron sequencing analysis of L1 retrotransposition and somatic mutation in the human brain. Cell. 2012;151(3):483–96.

8. Evrony GD, Lee E, Mehta BK, Benjamini Y, Johnson RM, Cai X, et al. Cell lineage analysis in human brain using endogenous retroelements. Neuron. 2015;85(1):49–59.

9. Muotri AR, Chu VT, Marchetto MC, Deng W, Moran JV, Gage FH. Somatic mosaicism in neuronal precursor cells mediated by L1 retrotransposition. Nature. 2005;435(7044):903–10.

10. Muotri AR, Zhao C, Marchetto MC, Gage FH. Environmental influence on L1 retrotransposons in the adult hippocampus. Hippocampus. 2009;19(10):1002–7.

11. Perrat PN, DasGupta S, Wang J, Theurkauf W, Weng Z, Rosbash M, et al. Transposition-driven genomic heterogeneity in the Drosophila brain. Science (New York, NY). 2013;340(6128):91–5.

12. Upton KR, Gerhardt DJ, Jesuadian JS, Richardson SR, Sanchez-Luque FJ, Bodea GO, et al. Ubiquitous L1 mosaicism in hippocampal neurons. Cell. 2015;161(2):228–39.

13. De Cecco M, Criscione SW, Peterson AL, Neretti N, Sedivy JM, Kreiling JA. Transposable elements become active and mobile in the genomes of aging mammalian somatic tissues. Aging. 2013;5(12):867–83.

14. Maxwell PH, Burhans WC, Curcio MJ. Retrotransposition is associated with genome instability during chronological aging. Proceedings of the National Academy of Sciences of the United States of America. 2011;108(51):20376–81.

15. Patterson MN, Scannapieco AE, Au PH, Dorsey S, Royer CA, Maxwell PH. Preferential retrotransposition in aging yeast mother cells is correlated with increased genome instability. DNA repair. 2015;34:18–27.

16. Savva YA, Jepson JE, Chang YJ, Whitaker R, Jones BC, St Laurent G, et al. RNA editing regulates transposon-mediated heterochromatic gene silencing. Nature communications. 2013;4:2745.

17. Wood JG, Hillenmeyer S, Lawrence C, Chang C, Hosier S, Lightfoot W, et al. Chromatin remodeling in the aging genome of Drosophila. Aging cell. 2010;9(6):971–8.

18. Wood JG, Helfand SL. Chromatin structure and transposable elements in organismal aging. Frontiers in genetics. 2013;4:274.

19. Li W, Prazak L, Chatterjee N, Gruninger S, Krug L, Theodorou D, et al. Activation of transposable elements during aging and neuronal decline in Drosophila. Nature neuroscience. 2013;16(5):529–31.

20. Kuo PH, Doudeva LG, Wang YT, Shen CK, Yuan HS. Structural insights into TDP-43 in nucleic-acid binding and domain interactions. Nucleic acids research. 2009;37(6):1799–808.

21. Ou SH, Wu F, Harrich D, Garcia-Martinez LF, Gaynor RB. Cloning and characterization of a novel cellular protein, TDP-43, that binds to human immunodeficiency virus type 1 TAR DNA sequence motifs. Journal of virology. 1995;69(6):3584–96.

22. Arai T, Hasegawa M, Akiyama H, Ikeda K, Nonaka T, Mori H, et al. TDP-43 is a component of ubiquitin-positive tau-negative inclusions in frontotemporal lobar degeneration and amyotrophic lateral sclerosis. Biochemical and biophysical research communications. 2006;351(3):602–11.

23. Neumann M. Molecular neuropathology of TDP-43 proteinopathies. International journal of molecular sciences. 2009;10(1):232–46.

24. Wang IF, Wu LS, Shen CK. TDP-43: an emerging new player in neurodegenerative diseases. Trends Mol Med. 2008;14(11):479–85.

25. Vanden Broeck L, Callaerts P, Dermaut B. TDP-43-mediated neurodegeneration: towards a loss-of-function hypothesis? Trends Mol Med. 2014;20(2):66–71.

26. Ling SC, Polymenidou M, Cleveland DW. Converging mechanisms in ALS and FTD: disrupted RNA and protein homeostasis. Neuron. 2013;79(3):416–38.

27. Chen-Plotkin AS, Lee VM, Trojanowski JQ. TAR DNA-binding protein 43 in neurodegenerative disease. Nature reviews Neurology. 2010;6(4):211–20.

28. Saberi S, Stauffer JE, Schulte DJ, Ravits J. Neuropathology of Amyotrophic Lateral Sclerosis and Its Variants. Neurol Clin. 2015;33(4):855–76.

29. Molliex A, Temirov J, Lee J, Coughlin M, Kanagaraj AP, Kim HJ, et al. Phase separation by low complexity domains promotes stress granule assembly and drives pathological fibrillization. Cell. 2015;163(1):123–33.

30. Lin Y, Protter DS, Rosen MK, Parker R. Formation and Maturation of Phase-Separated Liquid Droplets by RNA-Binding Proteins. Molecular cell. 2015;60(2):208–19.

31. Xiang S, Kato M, Wu LC, Lin Y, Ding M, Zhang Y, et al. The LC Domain of hnRNPA2 Adopts Similar Conformations in Hydrogel Polymers, Liquid-like Droplets, and Nuclei. Cell. 2015;163(4):829–39.

32. Liu-Yesucevitz L, Bilgutay A, Zhang YJ, Vanderweyde T, Citro A, Mehta T, et al. Tar DNA binding protein-43 (TDP-43) associates with stress granules: analysis of cultured cells and pathological brain tissue. PloS one. 2010;5(10):e13250.

33. Budini M, Buratti E. TDP-43 auto regulation: implications for disease. Journal of molecular neuroscience: MN. 2011;45(3):473–9.

34. Koyama A, Sugai A, Kato T, Ishihara T, Shiga A, Toyoshima Y, et al. Increased cytoplasmic TARDBP mRNA in affected spinal motor neurons in ALS caused by abnormal auto regulation of TDP-43. Nucleic acids research. 2016;44(12):5820–36.

35. Ciryam P, Kundra R, Morimoto RI, Dobson CM, Vendruscolo M. Supersaturation is a major driving force for protein aggregation in neurodegenerative diseases. Trends Pharmacol Sci. 2015;36(2):72–7.

36. Casci I, Pandey UB. A fruitful endeavor: modeling ALS in the fruit fly. Brain research. 2015;1607:47–74.

37. Gendron TF, Petrucelli L. Rodent models of TDP-43 proteinopathy: investigating the mechanisms of TDP-43-mediated neurodegeneration. Journal of molecular neuroscience: MN. 2011;45(3):486–99.

38. Janssens J, Van Broeckhoven C. Pathological mechanisms underlying TDP-43 driven neurodegeneration in FTLD-ALS spectrum disorders. Human molecular genetics. 2013;22(R1):R77–87.

39. Custer SK, Neumann M, Lu H, Wright AC, Taylor JP. Transgenic mice expressing mutant forms VCP/p97 recapitulate the full spectrum of IBMPFD including degeneration in muscle, brain and bone. Human molecular genetics. 2010;19(9):1741–55.

40. Ritson GP, Custer SK, Freibaum BD, Guinto JB, Geffel D, Moore J, et al. TDP-43 mediates degeneration in a novel Drosophila model of disease caused by mutations in VCP/p97. J Neurosci. 2010;30(22):7729–39.

41. Lanson NA, Jr., Maltare A, King H, Smith R, Kim JH, Taylor JP, et al. A Drosophila model of FUS-related neurodegeneration reveals genetic interaction between FUS and TDP-43. Human molecular genetics. 2011;20(13):2510–23.

42. He F, Krans A, Freibaum BD, Taylor JP, Todd PK. TDP-43 suppresses CGG repeat-induced neurotoxicity through interactions with HnRNP A2/B1. Human molecular genetics. 2014;23(19):5036–51.

43. Kawahara Y, Mieda-Sato A. TDP-43 promotes microRNA biogenesis as a component of the Drosha and Dicer complexes. Proceedings of the National Academy of Sciences of the United States of America. 2012;109(9):3347–52.

44. Ling JP, Pletnikova O, Troncoso JC, Wong PC. TDP-43 repression of nonconserved cryptic exons is compromised in ALS-FTD. Science (New York, NY). 2015;349(6248):650–5.

45. Emde A, Eitan C, Liou LL, Lib by RT, Rivkin N, Magen I, et al. Dysregulated miRNA biogenesis downstream of cellular stress and ALS-causing mutations: a new mechanism for ALS. The EMBO journal. 2015;34(21):2633–51.

46. Evrony GD, Lee E, Park PJ, Walsh CA. Resolving rates of mutation in the brain using single-neuron genomics. Elife. 2016;5.

47. Richardson SR, Morell S, Faulkner GJ. L1 retrotransposons and somatic mosaicism in the brain. Annu Rev Genet. 2014;48:1–27.

48. Douville R, Liu J, Rothstein J, Nath A. Identification of active loci of a human endogenous retrovirus in neurons of patients with amyotrophic lateral sclerosis. Annals of neurology. 2011;69(1):141–51.

49. Greenwood AD, Vincendeau M, Schmadicke AC, Montag J, Seifarth W, Motzkus D. Bovine spongiform encephalopathy infection alters endogenous retrovirus expression in distinct brain regions of cynomolgus macaques (Macaca fascicularis). Molecular neurodegeneration. 2011;6(11):44.

50. Kaneko H, Dridi S, Tarallo V, Gelfand BD, Fowler BJ, Cho WG, et al. DICER1 deficit induces Alu RNA toxicity in age-related macular degeneration. Nature. 2011;471(7338):325–30.

51. Lathe R, Harris A. Differential display detects host nucleic acid motifs altered in scrapie-infected brain. Journal of molecular biology. 2009;392(3):813–22.

52. Li W, Jin Y, Prazak L, Hammell M, Dubnau J. Transposable elements in TDP-43-mediated neurodegenerative disorders. PloS one. 2012;7(9):e44099.

53. Li W, Lee MH, Henderson L, Tyagi R, Bachani M, Steiner J, et al. Human endogenous retrovirus-K contributes to motor neuron disease. Sci Transl Med. 2015;7(307):307ra153.

54. Muotri AR, Marchetto MC, Coufal NG, Oefner R, Yeo G, Nakashima K, et al. L1 retrotransposition in neurons is modulated by MeCP2. Nature. 2010;468(7322):443–6.

55. Tan H, Qurashi A, Poidevin M, Nelson DL, Li H, Jin P. Retrotransposon activation contributes to fragile X premutation rCGG-mediated neurodegeneration. Human molecular genetics. 2012;21(1):57–65.

56. Alfahad T, Nath A. Retroviruses and amyotrophic lateral sclerosis. Antiviral Res. 2013;99(2):180–7.

57. MacGowan DJ, Scelsa SN, Imperato TE, Liu KN, Baron P, Polsky B. A controlled study of reverse transcriptase in serum and CSF of HIV-negative patients with ALS. Neurology. 2007;68(22):1944–6.

58. McCormick AL, Brown RH, Jr., Cudkowicz ME, Al-Chalabi A, Garson JA. Quantification of reverse transcriptase in ALS and elimination of a novel retroviral candidate. Neurology. 2008;70(4):278–83.

59. Steele AJ, Al-Chalabi A, Ferrante K, Cudkowicz ME, Brown RH, Jr., Garson JA. Detection of serum reverse transcriptase activity in patients with ALS and unaffected blood relatives. Neurology. 2005;64(3):454–8.

60. Crichton JH, Dunican DS, Maclennan M, Meehan RR, Adams IR. Defending the genome from the enemy within: mechanisms of retrotransposon suppression in the mouse germline. Cellular and molecular life sciences: CMLS. 2014;71(9):1581–605.

61. Malone CD, Hannon GJ. Small RNAs as guardians of the genome. Cell. 2009;136(4):656–68.

62. O’Donnell KA, Boeke JD. Mighty Piwis defend the germline against genome intruders. Cell. 2007;129(1):37–44.

63. Kabashi E, Lin L, Tradewell ML, Dion PA, Bercier V, Bourgouin P, et al. Gain and loss of function of ALS-related mutations of TARDBP (TDP-43) cause motor deficits in vivo. Human molecular genetics. 2010;19(4):671–83.

64. Ash PE, Zhang YJ, Roberts CM, Saldi T, Hutter H, Buratti E, et al. Neurotoxic effects of TDP-43 overexpression in C. elegans. Human molecular genetics. 2010;19(16):3206–18.

65. Romano M, Feiguin F, Buratti E. Drosophila Answers to TDP-43 Proteinopathies. J Amino Acids. 2012;2012:356081.

66. Miguel L, Frebourg T, Campion D, Lecourtois M. Both cytoplasmic and nuclear accumulations of the protein are neurotoxic in Drosophila models of TDP-43 proteinopathies. Neurobiology of disease. 2011;41(2):398–406.

67. Haidet-Phillips AM, Hester ME, Miranda CJ, Meyer K, Braun L, Frakes A, et al. Astrocytes from familial and sporadic ALS patients are toxic to motor neurons. Nature biotechnology. 2011;29(9):824–8.

68. Meyer K, Ferraiuolo L, Miranda CJ, Likhite S, McElroy S, Renusch S, et al. Direct conversion of patient fibroblasts demonstrates non-cell autonomous toxicity of astrocytes to motor neurons in familial and sporadic ALS. Proceedings of the National Academy of Sciences of the United States of America. 2014;111(2):829–32.

69. Chen H, Qian K, Chen W, Hu B, Blackbourn LWt, Du Z, et al. Human-derived neural progenitors functionally replace astrocytes in adult mice. The Journal of clinical investigation. 2015;125(3):1033–42.

70. Tong J, Huang C, Bi F, Wu Q, Huang B, Liu X, et al. Expression of ALS-linked TDP-43 mutant in astrocytes causes non-cell-autonomous motor neuron death in rats. The EMBO journal. 2013;32(13):1917–26.

71. Diaper DC, Adachi Y, Lazarou L, Greenstein M, Simoes FA, Di Domenico A, et al. Drosophila TDP-43 dysfunction in glia and muscle cells cause cytological and behavioural phenotypes that characterize ALS and FTLD. Human molecular genetics. 2013;22(19):3883–93.

72. Estes PS, Daniel SG, McCallum AP, Boehringer AV, Sukhina AS, Zwick RA, et al. Motor neurons and glia exhibit specific individualized responses to TDP-43 expression in a Drosophila model of amyotrophic lateral sclerosis. Disease models & mechanisms. 2013;6(3):721–33.

73. Romano G, Appocher C, Scorzeto M, Klima R, Baralle FE, Megighian A, et al. Glial TDP-43 regulates axon wrapping, GluRIIA clustering and fly motility by autonomous and non-autonomous mechanisms. Human molecular genetics. 2015.

74. Jin Y, Tam OH, Paniagua E, Hammell M. TEtranscripts: a package for including transposable elements in differential expression analysis of RNA-seq datasets. Bioinformatics. 2015;31(22):3593–9.

75. Prudencio M, Belzil VV, Batra R, Ross CA, Gendron TF, Pregent LJ, et al. Distinct brain transcriptome profiles in C9orf72-associated and sporadic ALS. Nature neuroscience. 2015;18(8):1175–82.

76. Song SU, Gerasimova T, Kurkulos M, Boeke JD, Corces VG. An env-like protein encoded by a Drosophila retroelement: evidence that gypsy is an infectious retrovirus. Genes & development. 1994;8(17):2046–57.

77. Ostertag EM, Prak ET, DeBerardinis RJ, Moran JV, Kazazian HH, Jr. Determination of L1 retrotransposition kinetics in cultured cells. Nucleic acids research. 2000;28(6):1418–23.

78. Estes PS, Boehringer A, Zwick R, Tang JE, Grigsby B, Zarnescu DC. Wild-type and A315T mutant TDP-43 exert differential neurotoxicity in a Drosophila model of ALS. Human molecular genetics. 2011;20(12):2308–21.

79. Hanson KA, Kim SH, Wassarman DA, Tibbetts RS. Ubiquilin modifies TDP-43 toxicity in a Drosophila model of amyotrophic lateral sclerosis (ALS). J Biol Chem. 2010;285(15):11068–72.

80. Li Y, Ray P, Rao EJ, Shi C, Guo W, Chen X, et al. A Drosophila model for TDP-43 proteinopathy. Proceedings of the National Academy of Sciences of the United States of America. 2010;107(7):3169–74.

81. Coyne AN, Yamada SB, Siddegowda BB, Estes PS, Zaepfel BL, Johannesmeyer JS, et al. Fragile X protein mitigates TDP-43 toxicity by remodeling RNA granules and restoring translation. Human molecular genetics. 2015;24(24):6886–98.

82. Chen Y, Pane A, Schupbach T. Cutoff and aubergine mutations result in retrotransposon upregulation and checkpoint activation in Drosophila. Current biology: CB. 2007;17(7):637–42.

83. Klattenhoff C, Bratu DP, McGinnis-Schultz N, Koppetsch BS, Cook HA, Theurkauf WE. Drosophila rasiRNA pathway mutations disrupt embryonic axis specification through activation of an ATR/Chk2 DNA damage response. Developmental cell. 2007; 12(1):45–55.

84. Brodsky MH, Weinert BT, Tsang G, Rong YS, McGinnis NM, Golic KG, et al. Drosophila melanogaster MNK/Chk2 and p53 regulate multiple DNA repair and apoptotic pathways following DNA damage. Molecular and cellular biology. 2004;24(3):1219–31.

85. Belgnaoui SM, Gosden RG, Semmes OJ, Haoudi A. Human LINE-1 retrotransposon induces DNA damage and apoptosis in cancer cells. Cancer cell international. 2006;6:13.

86. Aravin AA, Hannon GJ. Small RNA silencing pathways in germ and stem cells. Cold Spring Harbor symposia on quantitative biology. 2008;73:283–90.

87. Ghildiyal M, Zamore PD. Small silencing RNAs: an expanding universe. Nature reviews Genetics. 2009;10(2):94–108.

88. Saito K, Siomi MC. Small RNA-mediated quiescence of transposable elements in animals. Developmental cell. 2010;19(5):687–97.

89. Lee YS, Nakahara K, Pham JW, Kim K, He Z, Sontheimer EJ, et al. Distinct roles for Drosophila Dicer-1 and Dicer-2 in the siRNA/miRNA silencing pathways. Cell. 2004;117(1):69–81.

90. Czech B, Malone CD, Zhou R, Stark A, Schlingeheyde C, Dus M, et al. An endogenous small interfering RNA pathway in Drosophila. Nature. 2008;453(7196):798–802.

91. Zhou R, Hotta I, Denli AM, Hong P, Perrimon N, Hannon GJ. Comparative analysis of argonaute-dependent small RNA pathways in Drosophila. Molecular cell. 2008;32(4):592–9.

92. Vagin VV, Sigova A, Li C, Seitz H, Gvozdev V, Zamore PD. A distinct small RNA pathway silences selfish genetic elements in the germline. Science (New York, NY). 2006;313(5785):320–4.

93. Lippman Z, May B, Yordan C, Singer T, Martienssen R. Distinct mechanisms determine transposon inheritance and methylation via small interfering RNA and histone modification. PLoS biology. 2003;1(3):E67.

94. Sijen T, Plasterk RH. Transposon silencing in the Caenorhabditis elegans germ line by natural RNAi. Nature. 2003;426(6964):310–4.

95. Yang N, Kazazian HH, Jr. L1 retrotransposition is suppressed by endogenously encoded small interfering RNAs in human cultured cells. Nature structural & molecular biology. 2006;13(9):763–71.

96. Svoboda P, Stein P, Anger M, Bernstein E, Hannon GJ, Schultz RM. RNAi and expression of retrotransposons MuERV-L and IAP in preimplantation mouse embryos. Developmental biology. 2004;269(1):276–85.

97. Xie W, Donohue RC, Birchler JA. Quantitatively increased somatic transposition of transposable elements in Drosophila strains compromised for RNAi. PloS one. 2013;8(8):e72163.

98. Popova LM, Sakharova AV. [Virus-like inclusions in the cells of the central nervous system in amyotrophic lateral sclerosis]. Arkhiv patologii. 1982;44(10):28–33.

99. Saldi TK, Ash PE, Wilson G, Gonzales P, Garrido-Lecca A, Roberts CM, et al. TDP-1, the Caenorhabditis elegans ortholog of TDP-43, limits the accumulation of double-stranded RNA. The EMBO journal. 2014;33(24):2947–66.

100. Freibaum BD, Chitta RK, High AA, Taylor JP. Global analysis of TDP-43 interacting proteins reveals strong association with RNA splicing and translation machinery. Journal of proteome research. 2010;9(2):1104–20.

101. Peters L, Meister G. Argonaute proteins: mediators of RNA silencing. Molecular cell. 2007;26(5):611–23.

102. Robb GB, Rana TM. RNA helicase A interacts with RISC in human cells and functions in RISC loading. Molecular cell. 2007;26(4):523–37.

103. Pare JM, Tahbaz N, Lopez-Orozco J, LaPointe P, Lasko P, Hobman TC. Hsp90 regulates the function of argonaute 2 and its recruitment to stress granules and P-bodies. Molecular biology of the cell. 2009;20(14):3273–84.

104. Kim VN, Han J, Siomi MC. Biogenesis of small RNAs in animals. Nature reviews Molecular cell biology. 2009;10(2):126–39.

105. Dewannieux M, Esnault C, Heidmann T. LINE-mediated retrotransposition of marked Alu sequences. Nature genetics. 2003;35(1):41–8.

106. Tully T, Preat T, Boynton SC, Del Vecchio M. Genetic dissection of consolidated memory in Drosophila. Cell. 1994;79(1):35–47.

107. Qin H, Cressy M, Li W, Coravos JS, Izzi SA, Dubnau J. Gamma neurons mediate dopaminergic input during aversive olfactory memory formation in Drosophila. Current biology: CB. 2012;22(7):608–14.

108. Dietzl G, Chen D, Schnorrer F, Su KC, Barinova Y, Fellner M, et al. A genome-wide transgenic RNAi library for conditional gene inactivation in Drosophila. Nature. 2007;448(7150):151–6.

109. Rozhkov NV. Global Run-On Sequencing (GRO-seq) Library Preparation from Drosophila Ovaries. Methods Mol Biol. 2015;1328:217–30.

110. Dobin A, Davis CA, Schlesinger F, Drenkow J, Zaleski C, Jha S, et al. STAR: ultrafast universal RNA-seq aligner. Bioinformatics. 2013;29(1):15–21.

111. Langmead B, Trapnell C, Pop M, Salzberg SL. Ultrafast and memory-efficient alignment of short DNA sequences to the human genome. Genome Biol. 2009;10(3):R25.

112. Anders S, Huber W. Differential expression analysis for sequence count data. Genome Biol. 2010;11(10):R106.

113. Hu Y, Flockhart I, Vinayagam A, Bergwitz C, Berger B, Perrimon N, et al. An integrative approach to ortholog prediction for disease-focused and other functional studies. BMC Bioinformatics. 2011;12:357.

114. Ja WW, Carvalho GB, Mak EM, de la Rosa NN, Fang AY, Liong JC, et al. Prandiology of Drosophila and the CAFE assay. Proceedings of the National Academy of Sciences of the United States of America. 2007;104(20):8253–6.

115. Benzer S. BEHAVIORAL MUTANTS OF Drosophila ISOLATED BY COUNTERCURRENT DISTRIBUTION. Proceedings of the National Academy of Sciences of the United States of America. 1967;58(3):1112–9.

116. Chen G, Li W, Zhang QS, Regulski M, Sinha N, Barditch J, et al. Identification of synaptic targets of Drosophila pumilio. PLoS computational biology. 2008;4(2):e1000026.

117. McCloy RA, Rogers S, Caldon CE, Lorca T, Castro A, Burgess A. Partial inhibition of Cdk1 in G 2 phase overrides the SAC and decouples mitotic events. Cell cycle (Georgetown, Tex). 2014;13(9):1400–12.

118. Sahoo PK SS, and Wong AKC. A Survey of Thresholding Techniques. Computer Vision, Graphics, and Image Processing. 1988;41:233–60.

